# Spontaneous parthenogenesis in the parasitoid wasp *Cotesia typhae*: low frequency anomaly or evolving process?

**DOI:** 10.1101/2021.12.13.472356

**Authors:** Claire Capdevielle Dulac, Romain Benoist, Sarah Paquet, Paul-André Calatayud, Julius Obonyo, Laure Kaiser, Florence Mougel

## Abstract

Hymenopterans are haplodiploids and unlike most other Arthropods they do not possess sexual chromosomes. Sex determination typically happens via the ploidy of individuals: haploids become males and diploids become females. Arrhenotoky is believed to be the ancestral reproduction mode in Hymenopterans, with haploid males produced parthenogenetically, and diploid females produced sexually. However, a number of transitions towards thelytoky (diploid females produced parthenogenetically) have appeared in Hymenopterans, and in most cases populations or species are either totally arrhenotokous or totally thelytokous. Here we present the case of *Cotesia typhae* (Fernandez-Triana), a Braconidae that produces parthenogenetic females at a low frequency. The phenotyping of two laboratory strains and one natural population showed that this frequency is variable, and that this rare thelytokous phenomenon also happens in the wild. Moreover, mated females from one of the laboratory strains produce a few parthenogenetic daughters among a majority of sexual daughters. The analysis of daughters of heterozygous virgin females allowed us to show that a mechanism similar to automixis with central fusion is very likely at play in *C. typhae*. This mechanism allows some parts of the genome to remain heterozygous, especially at the chromosomes’ centromeres, which can be advantageous depending on the sex determination system involved. Lastly, in most species, the origin of thelytoky is either bacterial or genetic, and an antibiotic treatment as well as PCR experiments did not demonstrate a bacterial cause in *C. typhae*. The unusual case of low parthenogenetic frequency described in this species constitutes another example of the fascinating diversity of sex determination systems in Arthropods.

## Introduction

Sexual reproduction is the most widespread reproductive strategy among multicellular organisms and especially in animals. In contrast with its predominance, this reproductive mode appears costly due, for instance, to the necessity to detect and attract a partner, escape sexually transmitted diseases or avoid predation during mating. Because they share parenthood with their mate, sexual individuals transmit two-fold less of their genetic material to their progeny compared to asexual counterparts. The ubiquity of sex despite such disadvantages led to the definition of the so-called “paradox of sex” (Meirmans et al., 2012; Otto, 2009).

Numerous cases of evolution toward asexual reproduction or parthenogenesis have been reported, notably within arthropod taxa (The Tree of Sex Consortium, 2014). Parthenogenesis can produce either males (arrhenotoky) or females (thelytoky) from unfertilized eggs, but only the last case strictly coincides with asexual reproduction. It is also referred to as parthenogenesis *sensu stricto*. Thelytoky has been observed in almost all basal Hexapoda and non-holometabolous insect taxa (Vershinina and Kuznetsova, 2016) as well as in many holometabolous insect species (Gokhman and Kuznetsova, 2018). This wide taxonomic range illustrates the frequent transition from sexual to asexual taxa that arose independently in various lineages. This scattered distribution hides a global low percentage of parthenogenesis: thelytokous species represent less than 1% of the Hexapoda (Gokhman and Kuznetsova, 2018). The proportion of asexual lineages is also highly heterogeneous among taxa. Liegeois et al., (2021) detected frequencies between 0 and 6.7% among families of mayflies. Van der Kooi et al. (2017) reported frequencies ranging from 0 to 38% among genera of haplodiploid arthropods.

Transition from a sexual to asexual reproductive mode requires bypassing genetic and developmental constraints, a challenge that may be easier to face in some taxa than in others. In most species with a haplodiploid sex determination system, males develop from unfertilized eggs and are haploid while females develop from fertilized eggs leading to a diploid state. In such cases, embryonic development is initiated independently from egg fertilization and gametogenesis occurs through an abortive meiosis in haploid males (Ferree et al., 2019), both traits probably favouring an evolution toward thelytoky (Vorburger, 2014). The variable frequency of asexual reproduction even among haplodiploid lineages indicates that other factors allowing the transition toward this reproductive mode remain to be identified (van der Kooi et al., 2017).

The multiple and independent acquisitions of asexual reproduction are associated with numerous mechanisms that maintain or restore diploid state and produce females (Rabeling and Kronauer, 2013; Vorburger, 2014), as illustrated in Figure 1A. The figure focuses on genetic consequences in terms of heterozygosity, but each described situation may result from different cytological mechanisms. Clonal apomixis induces clonal reproduction and allows the complete preservation of heterozygosity. It may arise from mitosis, but also from endoreplication preceding meiosis with sister chromosome pairing, resulting in recombination between identical chromosomes (Archetti, 2010; Ma and Schwander, 2017). In automixis, meiosis occurs and is followed by different diploid restoration processes. Two meiosis products may assemble to generate a diploid cell: i) fusion of non-sister products separated during the first *reductional* division in central fusion or ii) fusion of sister cells produced during the second *equational* division in terminal fusion. Note that similar patterns are obtained when one meiotic division is suppressed to ensure the maintenance of a diploid state: the lack of first division is equivalent to central fusion while the lack of second division equates to terminal fusion. The restoration of diploidy may also result from gamete duplication involving either fusion of mitosis products or chromosomal replication without cellular division. In some lineages, the restoration of diploidy may operate during embryogenesis *via* endoreplication (Little et al., 2017; Pardo et al., 1995). Other mechanisms not illustrated here involve endoreplication followed by meiosis with non-sister chromosomes pairing or inverted meiosis with central or terminal fusion (Archetti, 2022, 2010). The consequences of thelytoky in terms of heterozygosity are variable depending on the mechanism: from complete homozygosity in one generation under gamete duplication to completely preserved heterozygosity in clonal apomixis, with intermediate levels of homozygosity in terminal and central fusion (Rabeling and Kronauer, 2013; Vorburger, 2014). According to the biology of a given species and the degree of necessity for maintaining heterozygosity, one or other mechanism may be favoured.

**Figure 1:**
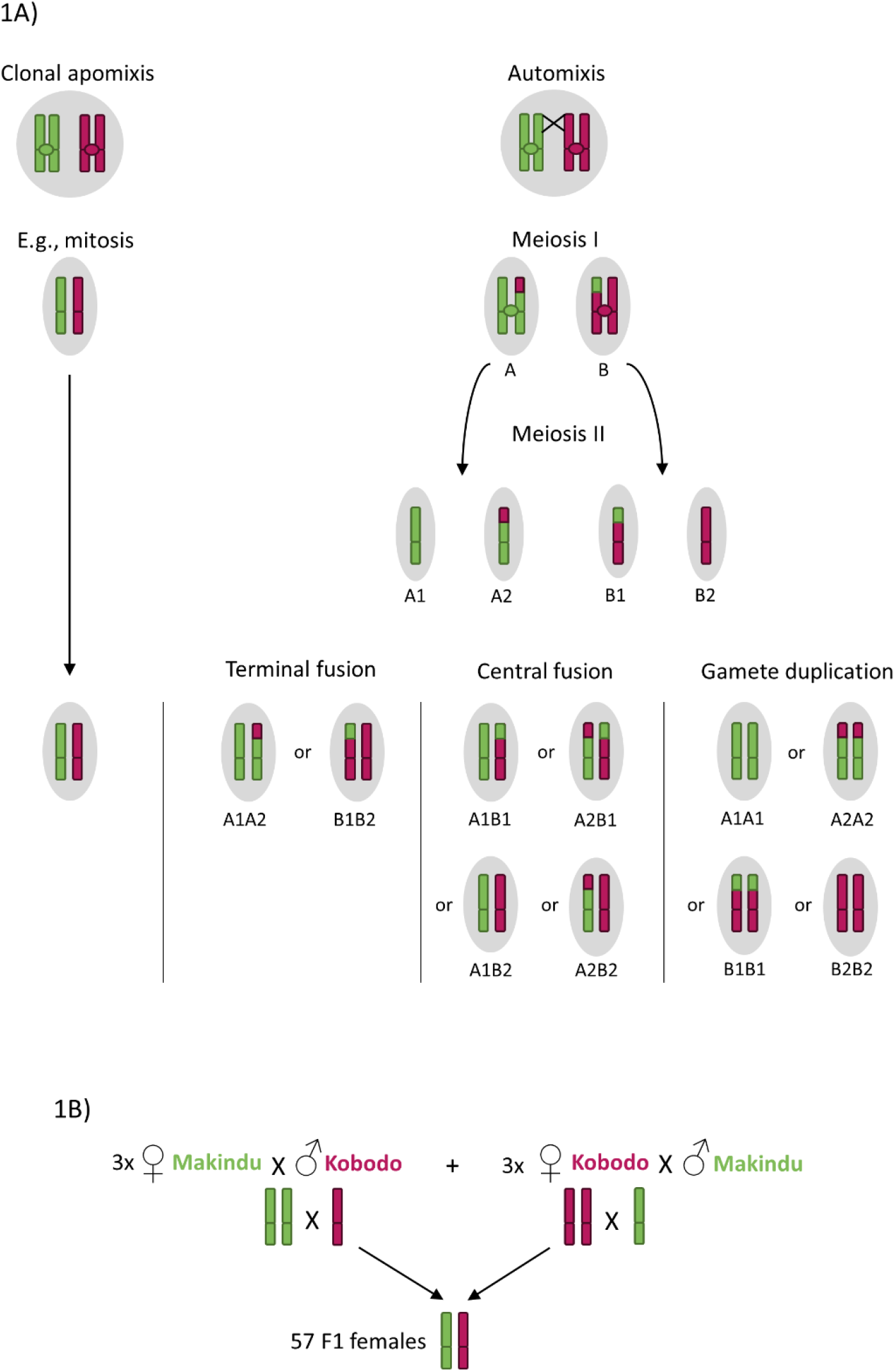
A) Expected genotype patterns of parthenogenetic daughters according to the main known thelytoky mechanisms. The cytological mechanisms are simplified because the figure focuses on the genetic consequences that can be achieved in different ways (see introduction for further details). The patterns can vary from complete loss of heterozygosity (under gamete duplication) to complete maintenance of heterozygosity (under clonal apomixis). B) Crosses performed in this study in order to obtain virgin heterozygous females. The analysis of recombination patterns of their daughters can give clues on the thelytoky mechanism involved.

Three main origins of thelytoky have been described: hybridization, bacterial endosymbiosis and genetic mutation (Tvedte et al., 2019). Hybridization, joining genomes from two distinct species, leads to improper chromosome pairing and dysfunctional meiosis that may promote asexuality (Morgan-Richards and Trewick, 2005). Endosymbiotic origin is the most widely studied cause of parthenogenesis (Ma and Schwander, 2017). To date, only bacteria have been shown to act as parthenogenesis inducers, but it is likely that other microorganisms could be involved. Most of the described causative agents belong to the genera *Wolbachia, Rickettsia* and *Cardinium*, endosymbionts also known to induce cytoplasmic incompatibility or feminization of male embryos. The particularity of endosymbiont induced parthenogenesis resides in its partial or total reversibility. Thelytokous species treated with antibiotics or heat may revert to sexual reproduction, although often performing less well than true sexual counterparts (Stouthamer et al., 1990). The genetic origin of thelytoky has often been suggested when antibiotic or heat treatment had no effect, but the precise identification of loci responsible for parthenogenesis has only been conducted in a few species (Chapman et al., 2015; Jarosch et al., 2011; Lattorff et al., 2005; Sandrock and Vorburger, 2011).

The relative frequency of thelytoky and sexual reproduction within species also varies according to taxa (Gokhman and Kuznetsova, 2018; Vershinina and Kuznetsova, 2016). Some species are described as obligate thelytokous when this mode of reproduction is the only one observed. Alternatively, thelytoky appears cyclic in some species where asexual generations alternate with sexual ones (Neiman et al., 2014). In other cases, polymorphism in the reproductive mode is observed either between populations (Foray et al., 2013; Leach et al., 2009) or within populations (Liu et al., 2019). Even in such polymorphic situations, thelytokous females usually produce female only progeny, albeit with a very low frequency of males in some cases, allowing rare events of sexual reproduction (Pijls et al., 1996).

Spontaneous occurrence of parthenogenesis has also been described in species reproducing via a sexual mode and qualified as tychoparthenogenesis (Ball, 2001; Pardo et al., 1995). Tychoparthenogenesis is characterized by a low hatching rate and a weak survival probability of the offspring (Little et al., 2017). It is typically considered as a dead-end accidental phenomenon in species adapted to sexual reproduction, although it may also correspond to an intermediate state in the evolution toward asexuality (van der Kooi and Schwander, 2015).

*Cotesia typhae* (Fernandez-Triana; Hymenoptera, Braconidae) is a gregarious endoparasitoid wasp native to Eastern Africa (Kaiser et al., 2017, 2015). It is specialized to one host, the corn stemborer *Sesamia nonagrioides* (Lefèbvre, Lepidoptera, Noctuidae). *Cotesia typhae* reproduces sexually and fertilized females typically lay 70-100 eggs in the first host encountered, among which about 70% develop into females and 30% into males (Benoist et al., 2020b). At least in laboratory conditions,sister-brother mating (sib-mating) frequently occurs indicating that inbreeding is not detrimental to this species. A genetic survey was conducted on this parasitoid wasp to compare two laboratory strains initiated from wild individuals sampled in two distant Kenyan localities, Kobodo and Makindu (Benoist et al., 2020a). The study led to the construction of a genetic map, based on crosses between the two strains. The phenotyping of the progenies obtained from these controlled crosses revealed an extremely variable sex-ratio, ranging from 100% to as low as 5% of females. Such a phenomenon could result from poorly mated females but also from rare thelytokous events in the progeny of unfertilized females. This last hypothesis was validated in a preliminary experiment allowing virgin females to oviposit. Among the numerous males emerging from the parasitized hosts, a few females were detected.

The aim of this study is to describe the low frequency thelytoky phenomenon in *Cotesia typhae* as a possible case of asexuality emergence. We first confirmed that this low frequency phenomenon is not restricted to artificial breeding but is also observed in natural conditions. We then tested whether thelytoky is induced by environmental conditions, that is lack of fertilisation, or observed in the progeny of mated as well as virgin females. Finally, we addressed the question of the mechanisms allowing the asexual production of females, because these mechanisms are tightly linked to the loss or conservation of heterozygosity and to the evolvability of asexual lineages.

## Methods

### Biological material

Two separate *Cotesia typhae* parasitoid strains were obtained from adults that emerged from naturally parasitized *Sesamia nonagrioides* caterpillars collected in the field at two localities in Kenya (Kobodo: 0.679S, 34.412E; West Kenya; 3 caterpillars collected in 2013 and Makindu: 2.278S, 37.825E; South-East Kenya; 10 caterpillars collected in 2010-2011). Isofemale lines were initiated in 2016 and inbred rearings have been subsequently kept for more than 80 generations at the Evolution, Génomes, Comportement et Ecologie laboratory (EGCE, Gif-sur-Yvette, France), where cross experiments and phenotyping were performed. The phenotyping of the wild population was performed on individuals that emerged from naturally parasitized *Sesamia nonagrioides* caterpillars collected in the field in 2020 at Kobodo (see above). The *S. nonagrioides* host strain used was initiated from caterpillars collected at Makindu (see above) and Kabaa (1.24S, 37.44E). The rearing protocol of *C. typhae* and *S. nonagrioides* is detailed in Benoist et al. (2020b).

### DNA extraction and genotyping methods

For our different experiments, DNA was extracted from *C. typhae* individuals using the NucleoSpin ® Tissue from Macherey Nagel, following the manufacturer’s instructions. For the experiment analysing the offspring of mated females, a direct PCR method was used instead of a classic DNA extraction because of the high number of individuals to be genotyped. In this case, for each individual, the abdomen was removed (because the presence of gametes could hinder the genotyping) and the rest of the body was placed in 20μL of Dilution Buffer and 0.5μL of DNA Release Additive (Thermoscientific). The tubes were kept at room temperature for 5 minutes then placed at 98°C for 2 minutes. One microliter of this mix or of DNA was then used as template for the following PCR.

Two different methods were used to genotype the chosen SNP markers, either HRM (High Resolution Melt), or allele specific PCR. HRM is based on the analysis of melt curves of DNA fragments after amplification by PCR. The melt curves are different according to the nucleotide composition of the DNA fragments, and therefore allow discrimination between homozygotes and heterozygotes at a given SNP. For each SNP marker, a 10μL mix was made with about 1ng of DNA, 0.2μM of each primer, and 5μL of Precision Melt Supermix (Bio-Rad), completed with water. The PCR protocol was 95°C for 2 minutes, followed by 40 cycles of 95°C for 10 seconds, 60°C for 30 seconds, 72°C for 30 seconds, followed by a complete denaturation of 30 seconds at 95°C before performing the melt curve. The melt curve was performed on a CFX96TM Real-Time System (Bio-Rad), and was started by an initial step of 1 minute at 60°C, followed by 10 seconds of every 0.2°C increment between 65°C and 95°C. The raw data resulting from the melt curves were analysed with the uAnalyze v2.1 software (Dwight et al., 2012) in order to infer the individuals’ genotypes at each SNP marker.

Two SNP markers (8225nov and 21770nov) were genotyped using allele specific PCR. For this method, two parallel amplifications were performed on individuals, one of the primers of the couple being a common primer, and the other one being a specific primer, either to the Makindu, or Kobodo allele. About 1ng of DNA was mixed with 1X buffer, 3mM MgCl2, 0.4mM dNTP, 0.4μM of each primer, 1U GoTaq® Flexi DNA Polymerase (Promega), and completed with water. The PCR programme was 5 minutes at 95°C, followed by 40 cycles of 95°C for 1 minute, 50°C or 55°C for 1 minute (50°C for 8225nov and 55°C for 21770nov), 72°C for one minute and a final elongation of 5 minutes at 72°C. The PCR products were then run on a 2% agarose gel to check which PCR were positive and therefore infer the genotypes.

All the primers used were designed for this study and their sequences are given in Supplementary Table 2.

### Phenotyping the strains/populations for the thelytokous character

In this study, the phenotyping consists of counting the number of males and females in the offspring of virgin females, to quantify the thelytoky phenomenon. To obtain virgin females, individual cocoons were isolated from cocoon masses and kept in tubes with a moistened cotton wool ball and a drop of honey at 27°C until the emergence of the adults. The virgin females were then each allowed to oviposit in one *S. nonagrioides* caterpillar, and the number of males and females in their offspring was counted after the development of the new *C. typhae* generation.

### Flow cytometry for ploidy analysis

Flow cytometry analysis was performed on one control female from a mixed cocoon mass (produced by a fertilized female), two control males, and five parthenogenetic females (produced by a virgin female), all coming from the Makindu laboratory strain, to determine their ploidy. The individuals were frozen in liquid nitrogen and processed in the Imagerie-Gif Platform of Institute for Integrative Biology of the Cell (I2BC), CNRS, Gif-sur-Yvette according to the protocol in (Bourge et al., 2018).

### Fecundity assessment of parthenogenetic females

Parthenogenetically produced females (hereafter referred as parthenogenetic females) were tested for their fecundity and their ability to perform thelytoky. To test whether mated parthenogenetic females had the same fecundity as mated control females, cocoon masses resulting from the eggs laid by *C. typhae* Makindu virgin females in *S. nonagrioides* caterpillars were divided in smaller cocoon packs to spot the few parthenogenetic females more easily among the males after the emergence of the adults. Adults were left together for one day with water and honey to allow mating, and eleven parthenogenetic females were then allowed to oviposit in S. nonagrioides caterpillars. After the emergence of the resulting offspring, the number of males and females was counted for each one of them and the sex-ratio was calculated for comparison with that obtained from fertilized females from the control Makindu laboratory strain. To test whether virgin parthenogenetic females were able to produce daughters parthenogenetically (these daughters will hereafter be referred as parthenogenetic daughters), all the cocoons from 5 virgin females’ progenies were isolated and the virgin parthenogenetic females emerging from these cocoons were allowed to oviposit in *S. nonagrioides* caterpillars. The number of males and females was then counted in each resulting offspring.

### Identifying the thelytoky mechanism in *Cotesia typhae*

To find out which thelytoky mechanism is at play in the *C. typhae* Makindu laboratory strain, virgin heterozygous females are needed to analyse the recombination patterns of their offspring. Indeed, according to the mechanism, the female offspring will be more or less heterozygous, as explained in introduction (Figure 1A). To obtain virgin heterozygous females, six controlled crosses were performed between the Makindu and Kobodo laboratory strains, 3 in each direction (Figure 1B). Prior to this, cocoons had been isolated from masses of each strain, in order to obtain virgin males and females for the crosses. Cocoons were then isolated from the masses resulting from the crosses, leading to the emergence of virgin F1 heterozygous females. 57 of these females were allowed to oviposit in S. nonagrioides caterpillars, and the offspring of the 57 resulting cocoon masses were sexed and counted. Six females and four males from the two parental strains (including the individuals used for the initial crosses), five F1 heterozygous females and the nine parthenogenetic females obtained through this experiment were kept for DNA extraction and genotyping, in order to analyse the recombination patterns resulting from parthenogenesis.

To analyse the recombination patterns of the nine parthenogenetic females, we genotyped 63 SNP (Single Nucleotide Polymorphism) markers, having different alleles between the Makindu and Kobodo strains, and being distributed along the 10 chromosomes of the genetic map of *Cotesia typhae*, (Benoist et al., 2020a). Four chromosomes contained more markers than the others to investigate the recombination patterns along chromosomes. For these four chromosomes, the markers were chosen to have about 10cM between two successive markers when possible. The genetic position of each marker is given in Supplementary Table 1, and the genotyping method used (HRM or allele specific PCR) is given in Supplementary Table 2.

**Table 1:** Results of the thelytoky phenotyping of the different strains/populations.

|  |  | N | Number of virgin females that produced female offspring (parthenogenetically) Percentage<sup>a</sup> [95% confidence interval] | Number of males produced by all virgin females | Number of females produced by all virgin females Percentage<sup>b</sup> [95% confidence interval] | Mean number [min ; max] of parthenogenetic females in the progenies containing females |
| --- | --- | --- | --- | --- | --- | --- |
| Makindu isofemale strain |  | 99 | 67<br>68% [57.4-76.5] | 10657 | 225<br>2% [1.81-2.36] | 3.4 [1;8] |
| Kobodo isofemale strain |  | 40 | 0 | 5405 | 0 | - |
| Kobodo wild population |  | 29 | 8<br>28% [13.4-47.5] | 2132 | 8<br>0.4% [0.17-0.77] | 1 [1;1] |
| Makindu strain after antibiotic treatment | After 4 generations | 55 | 15<br>27% [16.5-41.2] | 5886 | 17<br>0.3% [0.17-0.47] | 1.13 [1;2] |
|  | After 11 generations | 79 | 13<br>16% [9.4-26.9] | 8272 | 16<br>0.2% [0.11-0.32] | 1.23 [1;4] |

The phenotyping is based on the number and frequency of daughters produced parthenogenetically (parthenogenetic females) in the offspring of virgin *C. typhae* females. "N" is the total number of virgin females tested. a: percentage of all females tested. b: percentage of all offspring.

### Identifying the thelytoky phenomenon in mated females’ progenies

To see if mated females produce daughters parthenogenetically as well as sexually, 40 crosses between Makindu virgin females and Kobodo males were performed, according to the protocol described in the previous section. By genotyping the daughters of these crosses with a SNP marker differing between the Makindu and Kobodo strains, we can deduce if they were produced sexually (if heterozygous at the marker) or through parthenogenesis (if homozygous for the Makindu allele at the marker).

Out of the 40 crosses, five resulted in male only offspring and were removed from our analysis. The 35 remaining crosses lead to a mix of male and female offspring, for a total of 1861 daughters and 1803 sons. The 1861 daughters were genotyped at one SNP marker, 27068nov, according to the protocol described in the dedicated section (HRM technique). All the females that had a clear Makindu homozygote genotype and all the females presenting an uncertain genotype were then genotyped at 2 more markers (8225nov and 21770nov, allele specific PCR technique) to confirm their status.

### Search for a bacterial cause of thelytoky in *Cotesia typhae*

To find out if the cause of thelytoky in *C. typhae* could be bacterial, we first performed PCR with primers designed to amplify DNA sequences from several micro-organisms known to manipulate sex in insects. Ten virgin Makindu females that produced daughters parthenogenetically and two virgin Makindu females that didn’t produce daughters were tested with 8 primer sets, taken from Foray et al., (2013), except for one primer set, specific to *Wolbachia*, taken from Casiraghi et al., (2005). The primer sequences, Tm used for PCR, and their original publication are given in Supplementary Table 3. About 1ng of DNA was mixed with 1X buffer, 3mM MgCl2, O.4mM dNTP, O.4μM of each primer, 1U GoTaq® Flexi DNA Polymerase (Promega), and completed with water. The PCR programme was 5 minutes at 95°C, followed by 40 cycles of 95°C for 1 minute, Tm for 1 minute, 72°C for one minute and a final elongation of 5 minutes at 72°C. The PCR products were then run on a 1% agarose gel to check for positive amplification.

The amplified fragments obtained with the Arsenophonus primer set were sequenced with the BigDye™ Terminator v1.1 Cycle Sequencing Kit (ThermoFisher Scientific), following the manufacturer’s protocol. After the identification of the bacteria Pantoea dispersa through sequencing, 8 virgin Kobodo females that didn’t produce any parthenogenetic daughters were also tested with this primer set. Since our Kobodo laboratory strain doesn’t undergo thelytoky, this test was performed to check if *Pantoea dispersa* could be the causative agent of thelytoky in *C. typhae*.

To complete this experiment, we performed an antibiotic treatment on the Makindu laboratory strain to remove any potential sex manipulating bacteria in *C. typhae* females. Rifampicin was added in the host caterpillars’ artificial diet, at a final concentration of 2g/L, for 4 *C. typhae* generations. This treatment has previously been shown to eliminate *Wolbachia* bacteria in a close species, *Cotesia sesamiae* (Mochiah et al., 2002). The phenotyping results after this first treatment being ambiguous, it was continued for 7 more generations, with tetracyclin added for the 4 last generations, also at a final concentration of 2g/L. At that point, 79 virgin *C. typhae* females were phenotyped, according to the phenotyping protocol previously described.

## Results

### Phenotypes of the different strains/populations

We phenotyped two laboratory strains for the thelytokous character, Makindu and Kobodo, and a wild population, coming from the Kobodo locality. The numbers of available virgin females obtained for phenotyping were as follows: 99 from 13 different cocoon masses for the Makindu strain, 40 from 7 cocoon masses for the Kobodo strain, and 29 from 6 cocoon masses for the Kobodo wild population. The results (Table 1) are very contrasted between these three populations, since the number of parthenogenetically produced females (referred as parthenogenetic females) is null in the Kobodo isofemale strain, intermediate in the Kobodo wild population (28% of virgin females produced daughters), and high in the Makindu isofemale strain (68% of virgin females produced daughters). The thelytokous phenotype is therefore present in the wild and is not a laboratory artefact, but was apparently lost in the Kobodo laboratory strain (likely due to genetic drift), or present at a frequency too low to be detected. Unfortunately, the wild Makindu population does not exist anymore and could not be tested in this study.

### Ploidy of the daughters of virgin females

One daughter from the progeny of a fertilized female and 2 males, all belonging to the Makindu laboratory strain, were processed by flow cytometry as respective controls for diploid and haploid *Cotesia typhae* genomes. Five parthenogenetic daughters of virgin females were then processed, resulting in an estimated genome size identical to the control female and twice that of the control males (Table 2). We can therefore conclude that *C. typhae* parthenogenetic females are diploid and not the result of feminization of haploid eggs.

**Table 2:** Genome size estimated by flow cytometry.

| Sample | Size (pg) | Size (Mpb) | Ploidy |
| --- | --- | --- | --- |
| Control female | 0.47 | 458.27 | Diploid |
| Control male 1 | 0.26 | 251.84 | Haploid |
| Control male 2 | 0.25 | 241.12 | Haploid |
| Parthenogenetic female 1 | 0.48 | 467.51 | Diploid |
| Parthenogenetic female 2 | 0.48 | 473.24 | Diploid |
| Parthenogenetic female 3 | 0.49 | 475.28 | Diploid |
| Parthenogenetic female 4 | 0.49 | 475.91 | Diploid |
| Parthenogenetic female 5 | 0.5 | 484.26 | Diploid |

Parthenogenetic females have the same genome size as the control female, corresponding to about twice the males' haploid genome size.

### Fecundity of parthenogenetic females

Eleven parthenogenetic females (issued from virgin Makindu mothers) were randomly allowed to mate with their brothers and were used to parasitize eleven caterpillars. Out of these eleven females, 4 had male only offspring and 7 had a mixed offspring. The number of offspring per female and the sex-ratio are indicated in Table 3. No significant difference of the offspring size and sex ratio was observed between the control and parthenogenetic female datasets (p-value obtained following Mann-Whitney rank test was 0.559 for offspring number and 0.07 for sex-ratio). The fecundity of parthenogenetic females is therefore equivalent to that of the control females.

**Table 3:** Comparison of the fecundity between Makindu parthenogenetic females and control females of the same laboratory strain.

| | N | Mean number of offspring per mated female ± Standard Error | Mean female sex-ratio (when mated) ± Standard Error |
| --- | --- | --- | --- |
| Control Makindu strain<sup>a</sup> | 41 | 59 ± 4.2 | 0.78 ± 0.03 |
| Parthenogenetic females | 7 | 54.3 ± 9.2 | 0.65 ± 0.12 |

a: data from (Benoist et al., 2017). Mean female sex-ratio corresponds to the mean proportion of females among all offspring.

All the cocoons resulting from the egg-laying of five virgin females (385 cocoons in total) were isolated in order to obtain virgin parthenogenetic females. Fourteen females were thus obtained, out of which ten produced a progeny. Three out of these 10 progenies (30%) contained parthenogenetic females. As a result, a total of 6 females and 773 males was obtained across the ten progenies. Virgin parthenogenetic females are therefore able to produce parthenogenetic females themselves.

### Thelytoky mechanism occurring in *Cotesia typhae*

The genotyping of the 63 SNP markers first confirmed that fathers and mothers of the initial crosses between the Makindu and Kobodo laboratory strains were homozygotes for their strain’s alleles. The 57 virgin F1 daughters resulting from these crosses were thus heterozygous at the SNP markers, which was confirmed for the 5 F1 daughters that were genotyped. Each of these females successfully parasitized a host larva, and from the 57 resulting offspring, six contained parthenogenetic females (originating from 4 of the initial 6 crosses, 2 in each cross direction), corresponding to a total of 9 F2 parthenogenetic females and 6653 males. These 9 females were genotyped for the 63 SNPs. The genotypes and the deduced recombination events are presented in Supplementary Table 1. The recombination patterns of the 4 chromosomes genotyped with a higher density of markers are shown in Figure 2.

**Figure 2:**
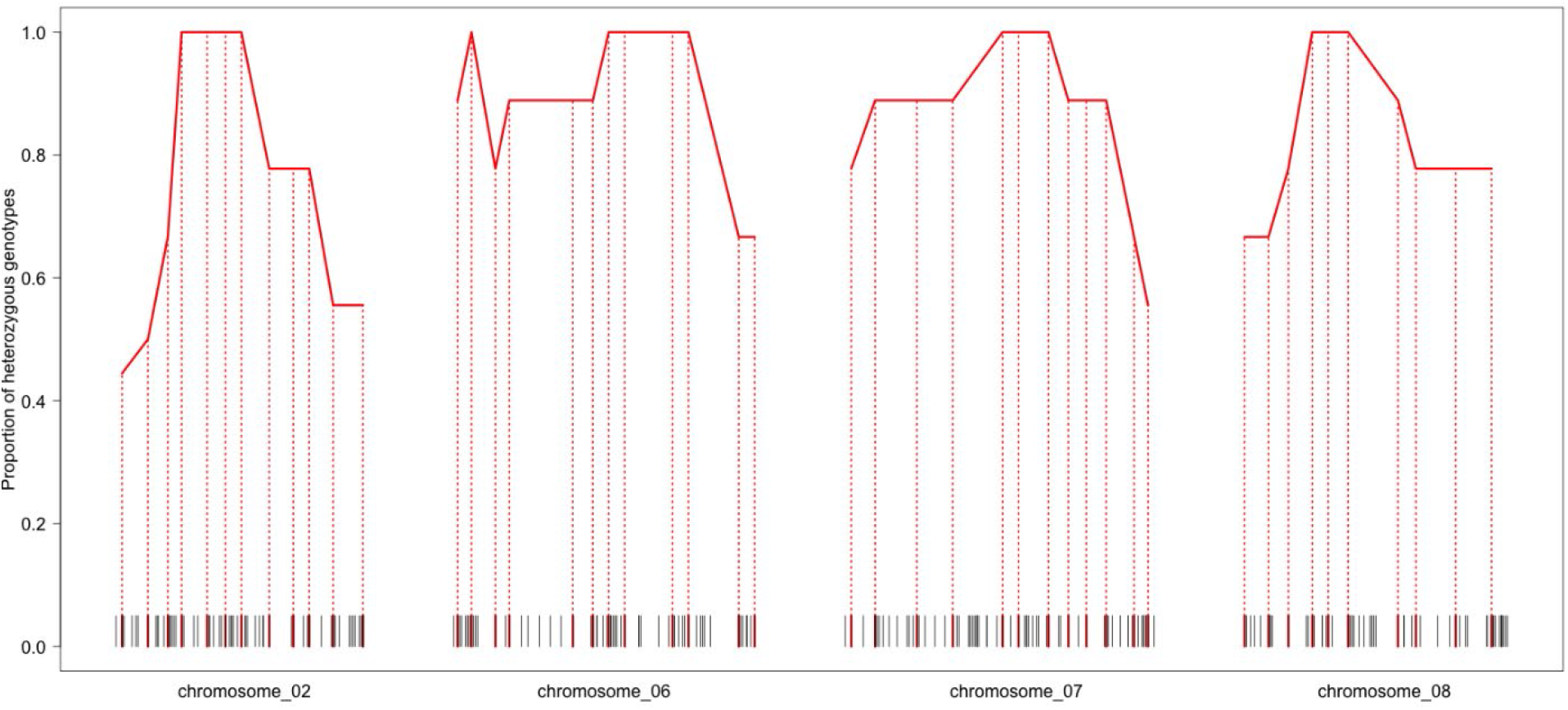
Proportion of heterozygous females (out of nine) for each genotyped SNP marker. The results are only shown for the 4 chromosomes for which a higher number of markers were genotyped. The black segments on the x axis are indicative of the genetic position of all the markers of the genetic map (Benoist et al., 2020a) and the red segments with the dotted lines correspond to the positions of the markers genotyped in this study. The occurrence of homozygous and heterozygous states along the chromosomes is congruent with a mechanism similar to automixis. Given the metacentric nature of *C. typhae* chromosomes (C. Bressac, personal communication), the observation of 100% heterozygosity in the central part of the chromosomes suggests that diploidy is either maintained by the suppression of the first meiotic division or restored through central fusion and is indicative of the position of each chromosome’s centromere.

For six of the females, a mixture of heterozygous and homozygous markers was observed, with a surplus of heterozygotes (280 heterozygous genotypes for 94 homozygous genotypes). The number and pattern of heterozygous markers for these females indicates a mechanism similar to automixis with central fusion (it could either result from the lack of first meiotic division or from the fusion of two non-sister meiotic products). Indeed, the central parts of the chromosomes maintain a heterozygous state while there is a recombination gradient leading to more homozygous genotypes towards the extremities of the chromosomes (Fig. 2). On average, nine recombination events per genome were detected for these six females with a minimum value of five events and a maximum of 16 events detected. Based on the density of the markers characterized, these results are consistent with the genetic length measured by Benoist et al. (2020a).

For the other 3 parthenogenetic females, all the 63 markers were heterozygous, revealing no detection of recombination event on the 10 chromosomes. We estimated the probability of such an observation under the hypothesis that a unique mechanism such as central fusion occurs. For each chromosome, we calculated a mean number of recombinations based on the nine parthenogenetic females. Assuming that the number of recombinations on a chromosome follows a Poisson distribution, we can estimate the probability of zero recombination for each chromosome based on the mean number estimate. It varies according to the genetic length of the chromosome and to the density of markers genotyped: it was estimated between 0.29 for chromosome 2 and 0.8 for chromosome 9. Multiplying the probability over the ten chromosomes, we calculated a probability of 0.0025 to observe an entirely heterozygous parthenogenetic daughter. Using this individual probability, we estimated that the probability to detect three out of nine parthenogenetic daughters showing no recombination events on the 10 chromosomes was 1.76 × 10^−6^. This hypothesis is very unlikely, therefore we suspect another mechanism could also be at play in causing thelytoky in *Cotesia typhae*. It is interesting to note that we observed both patterns (partial homozygosity and complete heterozygosity) in the offspring resulting from initial crosses of both directions. Moreover, one of the F1 heterozygous females displayed both patterns in her progeny. The raw data and the R script used to estimate the given probability are available at https://zenodo.org/record/6420801.

### Presence of parthenogenetic females among the daughters of mated females

Among the 40 crosses between Makindu females and Kobodo males, five gave male-only offspring and were excluded. The 35 remaining crosses that presented offspring comprising both males and females were kept in our analysis, leading to a total of 1861 females and 1803 males. All 1861 females were genotyped for one SNP marker. Females resulting from fertilization should be heterozygous while parthenogenetic females should be homozygous for the Makindu allele. In total, we found 14 homozygous females, which were confirmed by the genotyping of 2 other markers. These 14 females correspond to 0.77% of the parthenogenetic offspring (males resulting from arrhenotoky representing 99.33%) and originate from 10 different mothers (29% of the 35 mothers) (Table 4). Even though the percentage of parthenogenetic females found is much smaller than in the offspring of virgin females, this finding shows that the female progeny of mated females can come from a mixture of parthenogenesis and sexual reproduction.

**Table 4:**
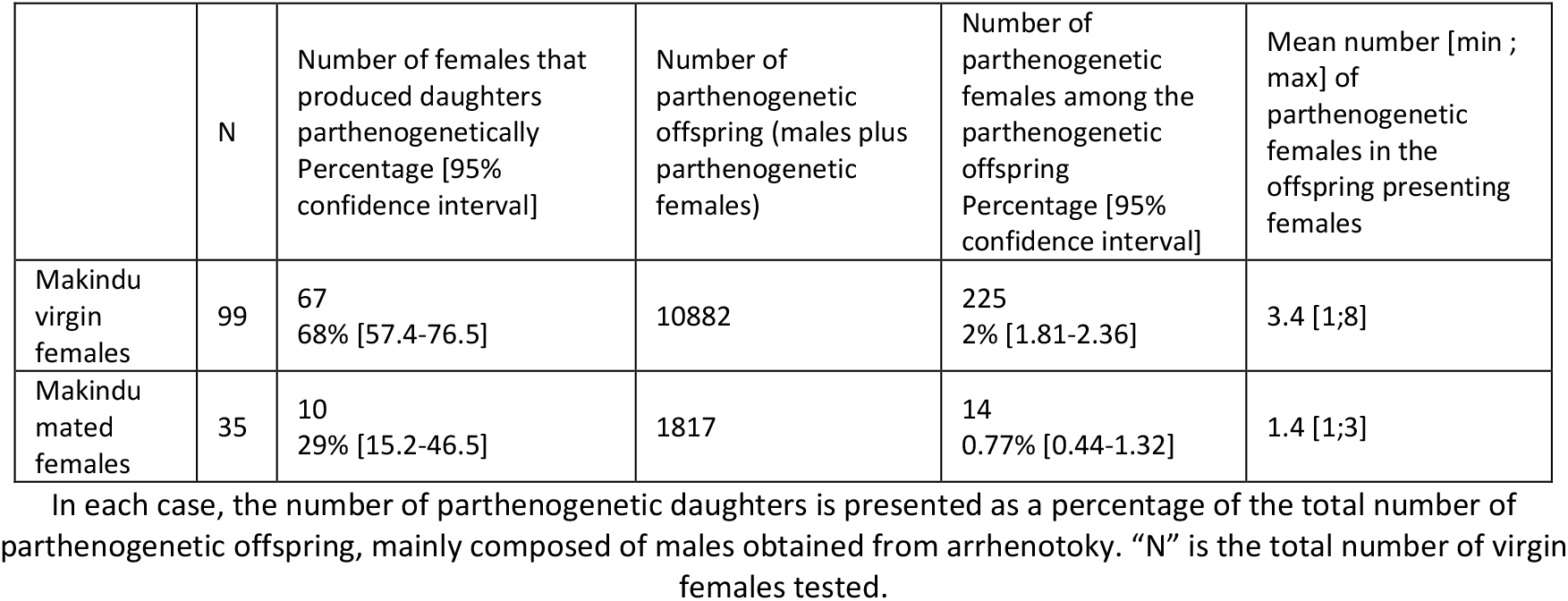
Comparison of the frequency of the thelytokous character between virgin and mated Makindu females.

|  | N | Number of females that produced daughters parthenogenetically<br>Percentage [95% confidence interval] | Number of parthenogenetic offspring (males plus parthenogenetic females) | Number of parthenogenetic females among the parthenogenetic offspring<br>Percentage [95% confidence interval] | Mean number [min ; max] of parthenogenetic females in the offspring presenting females |
| --- | --- | --- | --- | --- | --- |
| Makindu virgin females | 99 | 67<br>68% [57.4-76.5] | 10882 | 225<br>2% [1.81-2.36] | 3.4 [1;8] |
| Makindu mated females | 35 | 10<br>29% [15.2-46.5] | 1817 | 14<br>0.77% [0.44-1.32] | 1.4 [1;3] |

In each case, the number of parthenogenetic daughters is presented as a percentage of the total number of parthenogenetic offspring, mainly composed of males obtained from arrhenotoky. "N" is the total number of virgin females tested.

### Origin of thelytoky in *Cotesia typhae*

In order to find out if thelytoky in *Cotesia typhae* has a bacterial origin, we extracted DNA from virgin Makindu mothers that produced daughters and used primers to try to amplify the DNA of six different micro-organisms known for sex manipulation in insects: *Wolbachia, Ricketssia, Cardinium, Arsenophonus, Spiroplasma* and *Microsporidia* (Foray et al., 2013). Only one primer set led to a solid amplification, the one designed to amplify *Arsenophonus* 23S. After sequencing the amplified fragment, the bacterium was identified not as *Arsenophonus* but as *Pantoea dispersa*, for which no mention in relation to thelytoky was found in the literature. We then tried to amplify this same bacterium from the DNA of Kobodo virgin mothers, who don’t produce any daughters: *Pantoea dispersa* was present in all the samples tested. This makes it unlikely for this bacterium to be responsible for thelytoky in *C. typhae*.

After rearing parasitized caterpillars for four generations on a rifampicin diet, we phenotyped the Makindu strain again. 55 virgin females, coming from 5 different cocoon masses, were allowed to parasitize *Sesamia nonagrioides* caterpillars. Fifteen of these virgin females produced daughters, leading to a total of 17 daughters for 5886 sons (Table 1). Another phenotyping was performed on 79 females from 10 different cocoon masses, after 11 generations of a rifampicin diet (with tetracyclin added for the last 4 generations). Thirteen of these virgin females produced daughters, leading to a total of 16 daughters for 8272 sons (Table 1). The percentage of thelytokous females is thus smaller than the one observed before antibiotic treatment but non-zero.

### Discussion

The phenotypic survey presented here confirms the biological reality of low frequency asexual production of females in the haplo-diploid Hymenoptera *Cotesia typhae*. The process has been observed in a significant number of progenies from both an inbred laboratory strain and a natural population. It has been shown to occur in the progeny of virgin as well as fertilized females, despite concerning only a small fraction of the individuals from a cocoon mass.

This configuration of low frequency thelytoky is poorly illustrated in the literature, except in the well-studied species *Apis mellifera* for which the phenomenon has been described for a long time (Mackensen, 1943; Tucker, 1958). In the honey bee, both workers and virgin queens are able to produce a small proportion of females among unfertilized progeny (Gloag et al., 2019). However, in a common acceptation, the expression thelytoky rather refers to obligate parthenogenesis: it is defined as a “parthenogenetic mode where females produce only females from unfertilized eggs” (Vershinina and Kuznetsova, 2016). Among most illustrated examples of parthenogenesis *sensu stricto*, even when facultative, asexual production of females involves the whole progeny. The case described here is somewhat closer to what is called tychoparthenogenesis, based on the frequency of birth of parthenogenetic female eggs (Whiting, 1945). Tychoparthenogenesis is defined as “kind of occasional thelytoky characterized by the spontaneous hatching of a small proportion of eggs laid by virgin females” (Pardo et al., 1995). It has been mainly described in diplodiploid species where embryonic development is induced by sperm fertilization. In such species, developmental constraints and inbreeding depression prevent successful hatching of unfertilized eggs in most of the cases (Little et al., 2017). In haplodiploid species, unfertilized eggs hatch with a high frequency because they naturally produce males in species reproducing sexually. It is thus difficult to classify *C. typhae* as a tychoparthenogenetic species. Moreover, because the daughters produced parthenogenetically turned out to be viable and as fertile as sexually produced daughters in *C. typhae*, low frequency thelytoky appears to be either neutral or beneficial rather than disadvantageous as in most cases of tychoparthenogenesis described.

The question remains as to whether the phenomenon is accidental or an ongoing evolutionary process with an adaptive benefit (van der Kooi and Schwander, 2015). In the eusocial species *Apis mellifera*, low frequency thelytoky is clearly beneficial to workers when confronted with a queen-less colony (Gloag et al., 2019). This advantage led to an increased frequency of workers’ reproduction in a honey bee subspecies (*Apis m. capensis*) and to the development of social parasitism where egg-laying workers get adopted by a colony and compete the local queen (Neumann, 2001). Studying the occurrence of parthenogenesis among Ephemeroptera, Liegeois et al. (2021) suggested that asexual reproduction was selectively advantageous in many species from this insect order despite its associated low hatching success. The benefit derives from the short adult life and the low dispersal ability that reduce the probability of encountering a reproductive partner. The fitness of sexually reproducing individuals may consequently be reduced under certain circumstances. As in mayflies, *C. typhae* has an adult life limited to a few days (between two and three days in laboratory conditions, Kaiser et al., 2017). However, it is gregarious and mating between sisters and brothers emerging from the same cocoon mass is observed in rearing conditions. Female access to male fertilization should thus be facilitated unless sib-mating is avoided in natural conditions as demonstrated for its relative species *Cotesia glomerata* (Gu and Dorn, 2003) or *Venturia canescens* (Collet et al., 2020). Even in the presence of sexual partners, limited access to fertilization may also derive from ineffective copulation in cases of highly female biased sex ratios for example (Boivin, 2013). Whether it results from restricted access to males or to sperm itself, sperm limitation may favour expansion of asexual reproduction. Further experiments are needed to estimate mating and fertilization success of *Cotesia typhae* in natural conditions.

Beyond reproductive strategy itself, parthenogenesis has been shown to be associated with ecological characteristics that may favour or prevent its evolution. Two opposite ecological trends have been described co-occurring with asexual reproduction expansion: the “general purpose genotype” (GPG) where asexual lineages are observed on broader ecological niches than their sexual counterparts and the “frozen niche variation” (FNV) where parthenogenetic species or populations have far more restricted niches than sexual ones (Tvedte et al., 2019). Exploring a wide dataset of haplodiploid arthropods reproducing exclusively parthenogenetically (obligate parthenogenesis), van der Kooi et al. (2017) concluded that GPG was the most common situation. They showed that most parthenogenetic species have broader ecological and geographical range than close relative sexual species but also that transition toward parthenogenesis was more likely for species exhibiting a wide distribution. Studying the relative advantage of sexual and asexual reproduction, Song et al., (2012) developed a model based on resource availability, in spatial and temporal heterogeneous situations. They showed that sexual reproduction prevails in most of the cases but that asexual reproduction may be favoured when resource diversity is low and resource remains abundant over generations. Such a model corroborates numerous cases of ecological specialization of asexual lineages that have been described, such as *Venturia canescens*. In this polymorphic species, two kinds of populations live in sympatry: parthenogenetic populations found in stable anthropic habitats (bakeries and granaries) and sexual ones associated with natural and more instable resources (Schneider et al., 2002). Interestingly, before being characterized as a new species, *Cotesia typhae* was first identified as a specialized clade (only one host insect, *Sesamia nonagrioides*, mainly found on one host plant, *Typha domingiensis*) of the parasitoid species *Cotesia sesamiae*. According to (Branca et al., 2019), some populations of *C. sesamiae* are less specialized than others. Studying the existence of thelytokous reproduction in those populations would be informative about the possible link between emerging parthenogenesis and specialization.

Regarding the mechanism involved in thelytokous reproduction, we faced an unexpected result as data suggest that two different processes may co-occur: automixis with central fusion (or a similar cytological mechanism) and clonal apomixis. More surprisingly, the two supposed mechanisms were observed to cooccur in the progeny of a single female (K4M1) and independently of the cross direction to obtain F1 virgin mothers (Kobodo female x Makindu male or Makindu female x Kobodo male). Unfortunately, this result is supported by small sample size due to the scarcity of the phenomenon. We may wonder whether a unique mechanism, distinct from those already described, could explain such a result. Ma and Schwander (2017) describe for example an unusual process where meiosis is inverted (sister chromatids separate before homologs) followed by terminal fusion. However, the resulting progeny of such a process is 100% heterozygous, a result that does not differ from clonal apomixis. Another mechanism presented in the same review implies an endoreplication preceding meiosis. Assuming such a process occurs in *C. typhae*, and hypothesizing that recombination, and consequently segregation during the first division, may arise either between identical or between homologous chromosomes, some intermediate situations are expected. Once again, it does not reconcile the clear-cut figure we observe with individuals entirely heterozygous suggesting zero recombination between homologs and individuals for which recombination is observed for almost all homologs. If different mechanisms truly co-occur to produce females in an asexual way, it could reflect an accidental phenomenon arising from the relaxed control of sexual reproduction as observed in the honey bee (Aamidor et al., 2018). To better understand the mechanism underlying thelytoky in *C. typhae*, a cytological approach of meiosis and parthenogenesis would be necessary.

Despite the lack of a unified mechanism to explain the genotypic profile observed in the F2 progenies obtained, we can confirm that recombination occurred in at least 6 out of 9 cases, and that these recombination events were as frequent as those observed in sexual reproduction (Benoist et al., 2020a). By contrast, severe reductions of recombination rates were observed associated with parthenogenetic reproduction in the literature. For example, recombination in thelytokous workers is reduced by up to 10-fold in comparison to their sexually reproducing mothers in the Cape bee, *Apis mellifera capensis*, the social parasite of honeybee which reproduces parthenogenetically *via* automixis with central fusion (Baudry et al., 2004). In the little fire ant *Wasmannia auropunctata*, sexual populations coexist with asexual populations in which reproductive queens are produced by automictic parthenogenesis with central fusion. In asexual populations, recombination rate is reduced by a factor of 45 compared to the sexual populations (Rey et al., 2011). The reduction of recombination rate is assumed to mitigate the potential deleterious impact of thelytoky: under automixis with central fusion, heterozygosity is preserved unless recombination occurs (Figure 1). In species affected by inbreeding depression, a homozygosity increase would be detrimental and could be advantageously limited by a low recombination rate. As the molecular mechanisms involved in thelytoky and recombination are probably distinct, the situation observed in the Cape bee and little fire ant may result from a long-term evolutionary process. If the phenomenon described in *C. typhae* is recent, it may explain the unchanged recombination rate.

Another explanation could be that the inbreeding impact is meaningless in *C. typhae*. Hymenoptera are haplodiploid and could thus be less sensitive to inbreeding because most of the deleterious alleles are purged at the haploid state in males (Hedrick and Parker, 1997; Henter, 2003). However, their sex determination system may be highly compelling regarding homozygosity and ability to reproduce *via* thelytoky (Vorburger, 2014). The most common, and likely ancestral, sex determination system is governed by the genotype at one (sl-CSD: single locus Complementary Sex Determination) or few loci (ml-CSD: multi locus CSD) (Heimpel and de Boer, 2008). Under such a determinism, individuals that are heterozygous at least at one of these CSD loci develop as diploid females while hemizygous or homozygous individuals at all CSD loci develop as haploid or diploid males respectively. In most hymenopteran species, diploid males have a low survival rate and/or are often sterile. Enhanced homozygosity due to thelytoky may be very costly when it results in diploid male production (de Boer et al., 2015, 2012; van Wilgenburg et al., 2006; Zhou et al., 2007). Other sex determination systems that are less sensitive to homozygosity have been described in Hymenoptera, such as Paternal Genome Elimination (Heimpel and de Boer, 2008) or genome imprinting, described in *Nasonia vitripennis* (van de Zande and Verhulst, 2014). The mechanism of sex determination is unknown in *C. typhae*. However, it has been reared for more than 80 generations in laboratory conditions starting from an isofemale line and seems poorly sensitive to homozygosity increase.

The bacterial origin of thelytoky in *C. typhae* could not be either confirmed or completely discarded in the present study as an intermediate state (in terms of frequency of parthenogenesis) was observed following antibiotic treatment. The knowledge of the genetic mechanism could give some clues about the origin of parthenogenesis as endosymbionts have been mainly shown to favour gamete duplication. However, detailing specific interactions reveals a more complex picture. *Cardinium* is able to feminize diploid males (Giorgini et al., 2009) but also to induce automixis with central fusion (Zchori-Fein and Perlman, 2004). *Rickettsia* and *Wolbachia* have been shown to trigger clonal apomixis (Adachi-Hagimori et al., 2008; Weeks et al., 2011), although *Wolbachia* is mainly known to induce gamete duplication (Leach et al., 2009; Ma and Schwander, 2017). Furthermore, the list of endosymbionts is probably partial and in most of the documented examples of parthenogenesis endosymbiotically determined, the cytological mechanism remains unknown. Evidence that microorganisms can promote all processes of parthenogenesis will probably arise from future research. The examples of demonstrated genetic determinism of thelytoky are rare and only concern automixis with central fusion. This is the case for the Cape honey bee (Verma and Ruttner, 1983) and for the wasp *Venturia canescens* (Beukeboom and Pijnacker, 2000). However, Tsutsui et al., (2014) described a clonal apomixis mechanism in the parasitoid wasp *Meteorus pulchricornis* for which they proposed a genetic origin of thelytoky. Even more than for endosymbiont origin, genetic determinism of parthenogenesis requires thorough investigations to determine whether it is restricted to a few cytological mechanisms. Anyway, the clearly evidenced mechanism of autoximis with central fusion in *C. typhae* does not allow to settle between genetic and endosymbiont origin as this mechanism is common to both situations.

### Conclusion

In this study, we described an unusual example of low frequency thelytokous reproduction within a sexually reproducing species. We demonstrated that this asexual reproduction is likely the result of different mechanisms and occurs even in the progeny of fertilized females, an undescribed phenomenon to our knowledge. As most studies on asexual reproduction focus on obligate or cyclical parthenogenesis, we may wonder whether such low frequency and probably accidental thelytoky is common but until now mostly undetected among sexually reproductive species. It is well known that asexuality has emerged many times in numerous lineages. If the occurrence of accidental parthenogenesis turns out to be usual, it could represent the first step of evolutionary trajectories favoured either by reproductive or ecological advantages of asexual lineages.

## Acknowledgements

The present work has benefited from the I2BC Cytometry platform (Université Paris-Saclay, CEA, CNRS, Institute for Integrative Biology of the Cell (I2BC), 91198, Gif-sur-Yvette, France) with the help of Micka ёl Bourge. We thank Rémi Jeannette and Sylvie Nortier for insect rearing at Gif, and Daniel Couch for proofreading.

All experimentations were realized under the juridical frame of a Material Transfer Agreement signed between IRD, icipe and CNRS (CNRS 072057/IRD 302227/00) and the authorization to import *Cotesia* in France delivered by the DRIAAF of Ile de France (IDF 2017-OI-26-032).

We thank Christoph Haag, Jens Bast and Michael Lattorff for their constructive comments that helped us improve this paper.

Version 6 of this preprint has been peer-reviewed and recommended by Peer Community In Evolutionary Biology (https://doi.org/10.24072/pci.evolbiol.100141).

## Data, scripts and codes availability

Raw data for Figure 2, and Tables 1, 3 and 4 are available at https://zenodo.org/record/6420801

---

**Supplementary Table 1:**
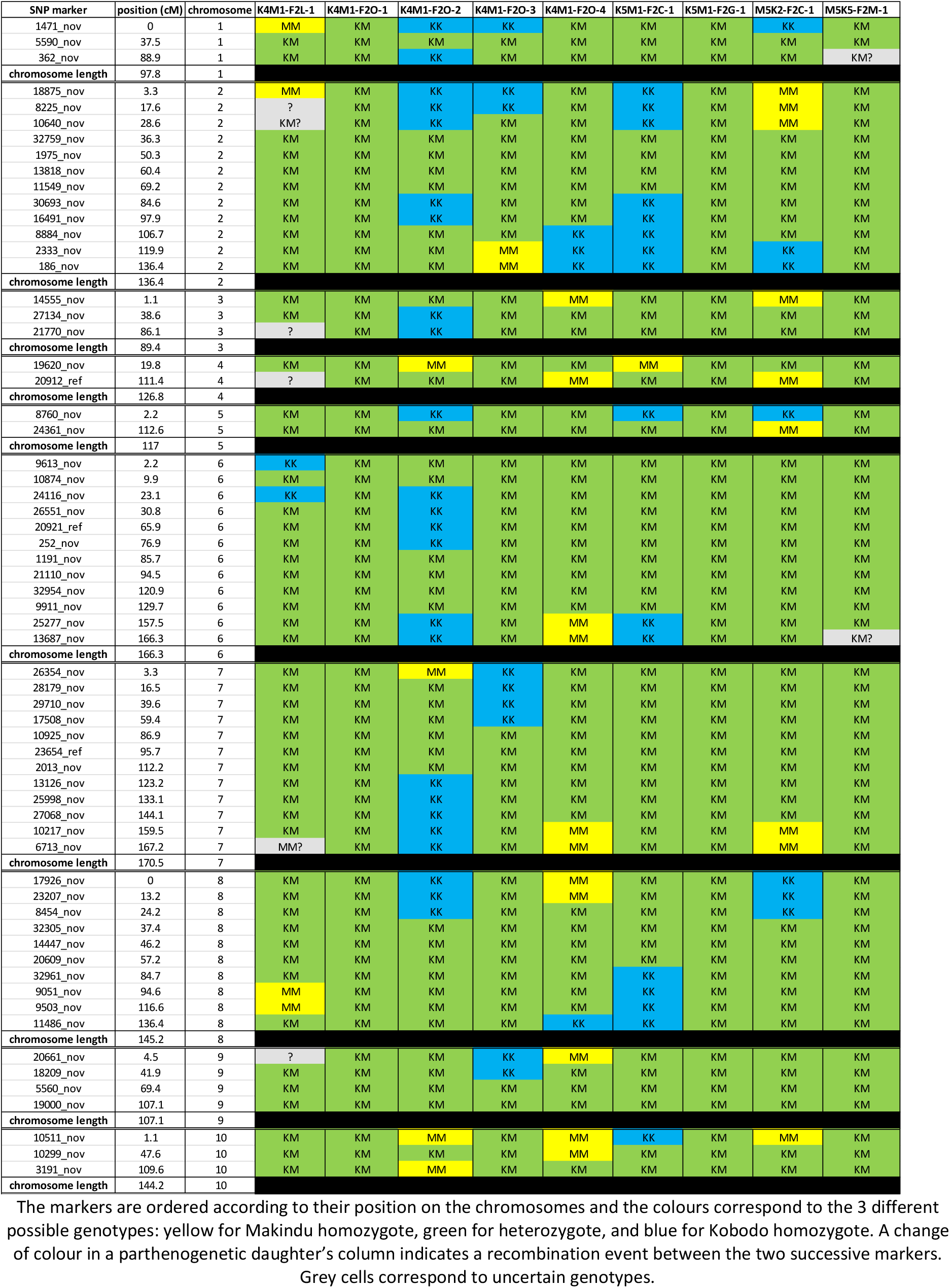
Genotypes of the 9 parthenogenetic daughters of virgin heterozygous females at all the 63 SNP markers used.

**Supplementary Table 2:**
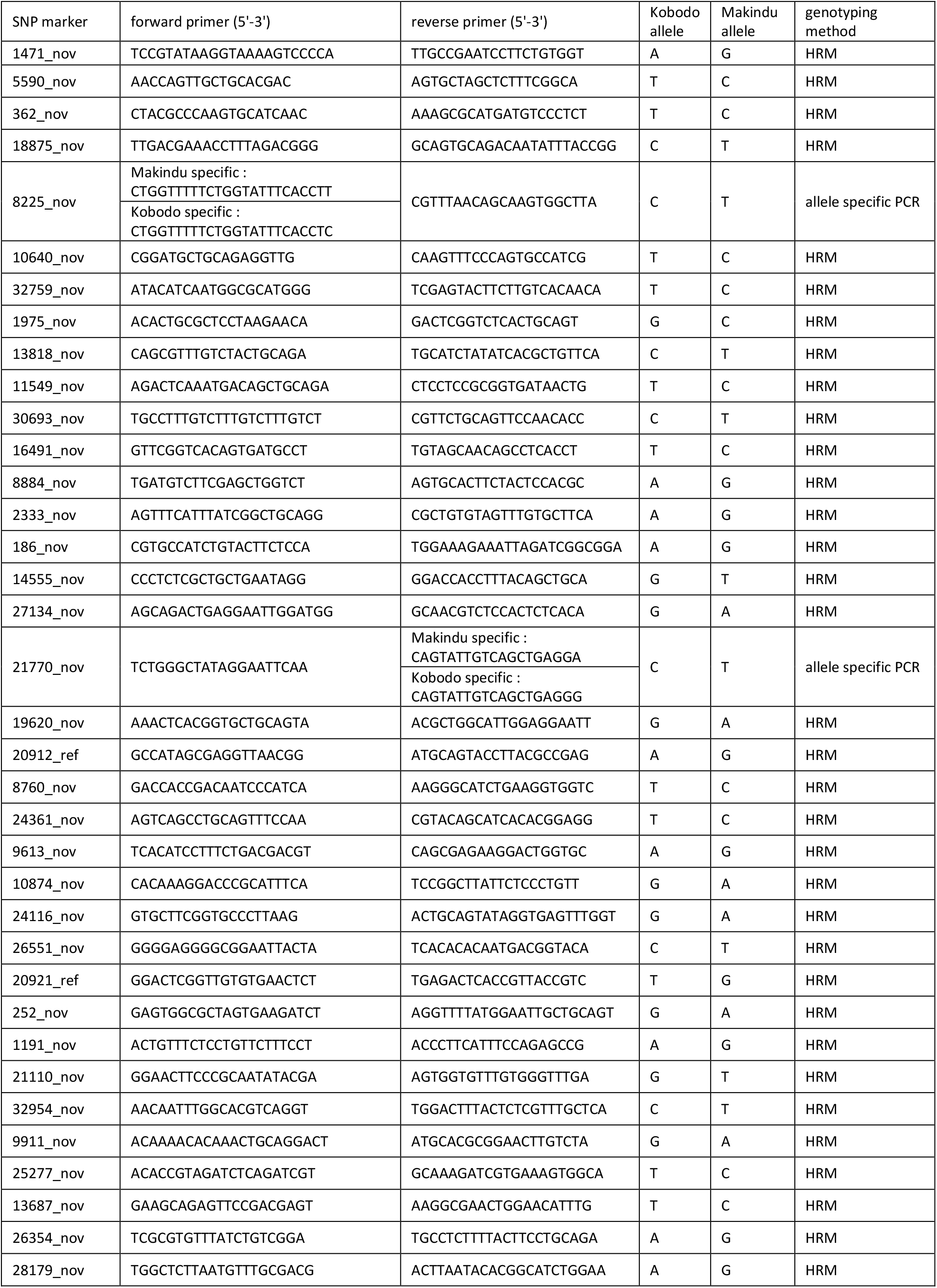
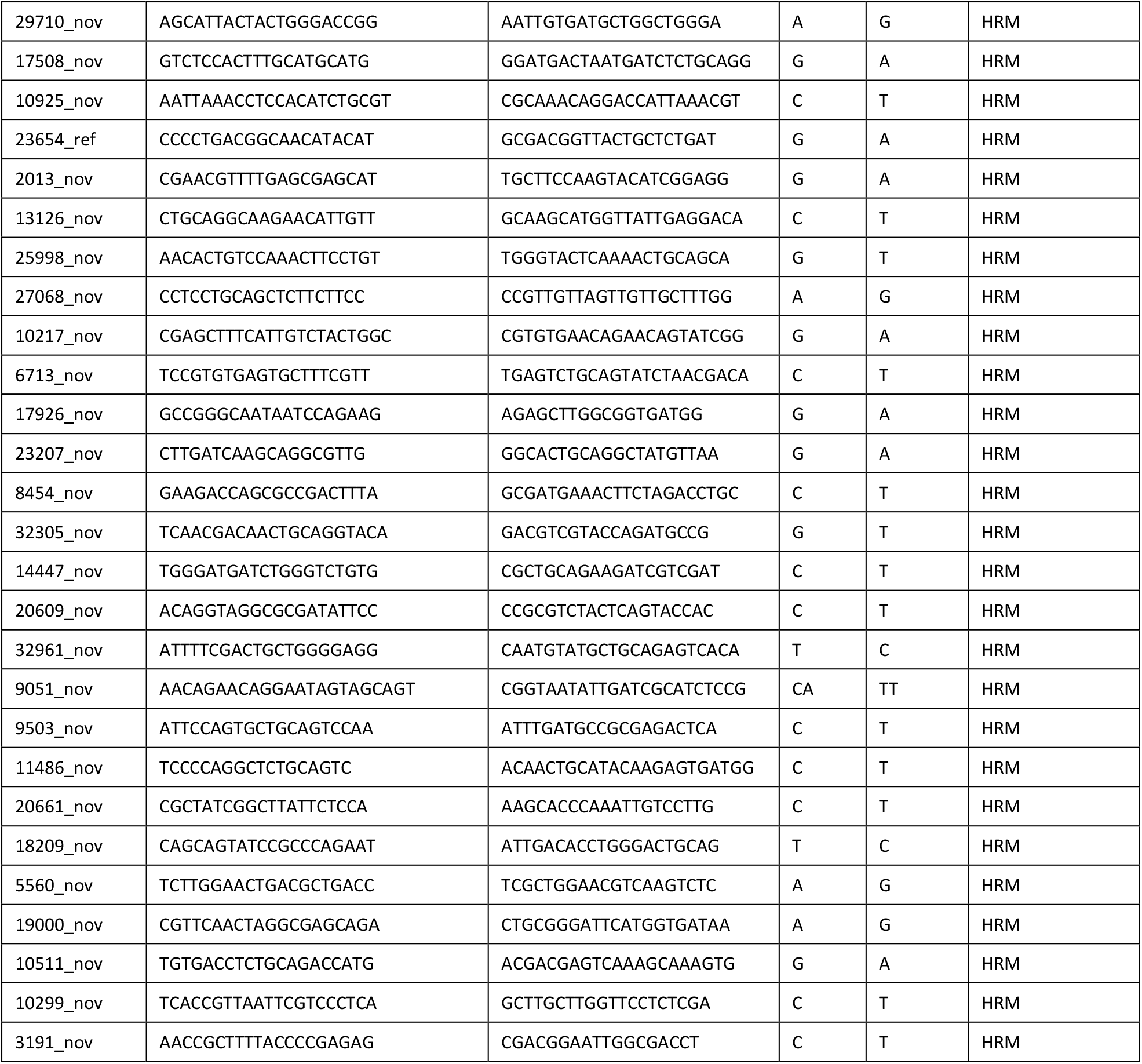
Primers’ sequences for the 63 SNP markers used for genotyping. The Kobodo and Makindu reference alleles are also indicated for each SNP.

**Supplementary Table 3:**
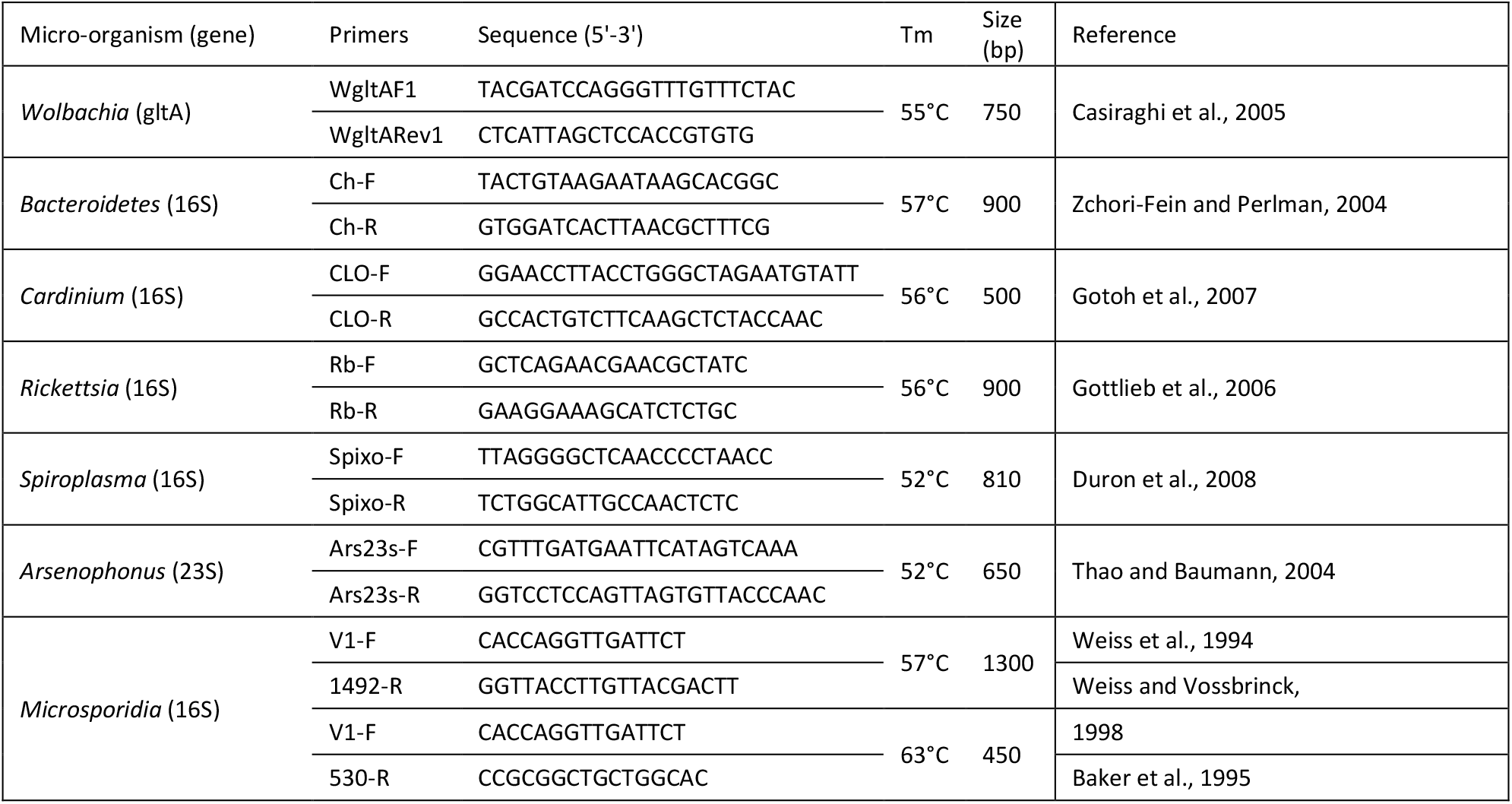
List of the micro-organisms whose presence was tested in *C. typhae* Makindu females, by PCR with specific primers.

## References

Aamidor, S.E., Yagound, B., Ronai, I., Oldroyd, B.P., 2018. Sex mosaics in the honeybee: how haplodiploidy makes possible the evolution of novel forms of reproduction in social Hymenoptera. Biol. Lett. 14, 20180670. https://doi.org/10.1098/rsbl.2018.0670

Adachi-Hagimori, T., Miura, K., Stouthamer, R., 2008. A new cytogenetic mechanism for bacterial endosymbiont-induced parthenogenesis in Hymenoptera. Proc. R. Soc. B. 275, 2667–2673. https://doi.org/10.1098/rspb.2008.0792

Archetti, M., 2022. Evidence from automixis with inverted meiosis for the maintenance of sex by loss of complementation. J of Evolutionary Biology 35, 40–50. https://doi.org/10.1111/jeb.13975

Archetti, M., 2010. Complementation, Genetic Conflict, and the Evolution of Sex and Recombination. Journal of Heredity 101, S21–S33. https://doi.org/10.1093/jhered/esq009

Ball, S.L., 2001. Tychoparthenogenesis and mixed mating in natural populations of the mayfly *Stenonema femoratum*. Heredity 87, 373–380. https://doi.org/10.1046/j.1365-2540.2001.00930.x

Baudry, E., Kryger, P., Allsopp, M., Koeniger, N., Vautrin, D., Mougel, F., Cornuet, J.M., Solignac, M., 2004. Whole-Genome Scan in Thelytokous-Laying Workers of the Cape Honeybee (*Apis mellifera capensis):* Central Fusion, Reduced Recombination Rates and Centromere Mapping Using Half-Tetrad Analysis. Genetics 252, 243–252. https://doi.org/10.1534/genetics.167.1.243

Benoist, R., Capdevielle-Dulac, C., Chantre, C., Jeannette, R., Calatayud, P., Drezen, J., Dupas, S., Le Rouzic, A., Le Ru, B., Moreau, L., Van Dijk, E., Kaiser, L., Mougel, F., 2020a. Quantitative trait loci involved in the reproductive success of a parasitoid wasp. Mol Ecol 29, 3476–3493. https://doi.org/10.1111/mec.15567

Benoist, R., Chantre, C., Capdevielle-Dulac, C., Bodet, M., Mougel, F., Calatayud, P.A., Dupas, S., Huguet, E., Jeannette, R., Obonyo, J., Odorico, C., Silvain, J.F., Le Ru, B., Kaiser, L., 2017. Relationship between oviposition, virulence gene expression and parasitism success in *Cotesia typhae nov. sp*. parasitoid strains. Genetica 145, 469–479. https://doi.org/10.1007/s10709-017-9987-5

Benoist, R., Paquet, S., Decourcelle, F., Guez, J., Jeannette, R., Calatayud, P.-A., Le Ru, B., Mougel, F., Kaiser, L., 2020b. Role of egg-laying behavior, virulence and local adaptation in a parasitoid’s chances of reproducing in a new host. Journal of Insect Physiology 120, 103987. https://doi.org/10.1016/j.jinsphys.2019.103987

Beukeboom, L.W., Pijnacker, L.P., 2000. Automictic parthenogenesis in the parasitoid Venturia canescens (Hymenoptera: Ichneumonidae) revisited 43, 6. https://doi.org/10.1139/g00-061

Boivin, G., 2013. Sperm as a limiting factor in mating success in Hymenoptera parasitoids. Entomol Exp Appl 146, 149–155. https://doi.org/10.1111/j.1570-7458.2012.01291.x

Bourge, M., Brown, S.C., Siljak-Yakovlev, S., 2018. Flow cytometry as tool in plant sciences, with emphasis on genome size and ploidy level assessment. Genetics & Applications 2, 1–12. https://doi.org/10.31383/ga.vol2iss2pp1-12

Branca, A., Le Ru, B., Calatayud, P.-A., Obonyo, J., Musyoka, B., Capdevielle-Dulac, C., Kaiser-Arnauld, L., Silvain, J.-F., Gauthier, J., Paillusson, C., Gayral, P., Herniou, E.A., Dupas, S., 2019. Relative Influence of Host, *Wolbachia*, Geography and Climate on the Genetic Structure of the Sub-saharan Parasitic Wasp *Cotesia sesamiae*. Front. Ecol. Evol. 7, 309. https://doi.org/10.3389/fevo.2019.00309

Casiraghi, M., Bordenstein, S.R., Baldo, L., Lo, N., Beninati, T., Wernegreen, J.J., Werren, J.H., Bandi, C., 2005. Phylogeny of *Wolbachia pipientis* based on gltA, groEL and ftsZ gene sequences: clustering of arthropod and nematode symbionts in the F supergroup, and evidence for further diversity in the *Wolbachia* tree. Microbiology 151, 4015–4022. https://doi.org/10.1099/mic.0.28313-0

Chapman, N.C., Beekman, M., Allsopp, M.H., Rinderer, T.E., Lim, J., Oxley, P.R., Oldroyd, B.P., 2015. Inheritance of thelytoky in the honey bee *Apis mellifera capensis*. Heredity 114, 584–592. https://doi.org/10.1038/hdy.2014.127

Collet, M., Amat, I., Sauzet, S., Auguste, A., Fauvergue, X., Mouton, L., Desouhant, E., 2020. Insects and incest: Sib-mating tolerance in natural populations of a parasitoid wasp. Mol Ecol 29, 596–609. https://doi.org/10.1111/mec.15340

de Boer, J.G., Groenen, M.A., Pannebakker, B.A., Beukeboom, L.W., Kraus, R.H., 2015. Population-level consequences of complementary sex determination in a solitary parasitoid. BMC Evol Biol 15, 98. https://doi.org/10.1186/s12862-015-0340-2

de Boer, J.G., Kuijper, B., Heimpel, G.E., Beukeboom, L.W., 2012. Sex determination meltdown upon biological control introduction of the parasitoid *Cotesia rubecula?* Evol Appl 5, 444–454. https://doi.org/10.1111/j.1752-4571.2012.00270.x

Dwight, Z.L., Palais, R., Wittwer, C.T., 2012. uAnalyze: Web-Based High-Resolution DNA Melting Analysis with Comparison to Thermodynamic Predictions. IEEE/ACM Trans. Comput. Biol. and Bioinf. 9, 1805–1811. https://doi.org/10.1109/TCBB.2012.112

Ferree, P.M., Aldrich, J.C., Jing, X.A., Norwood, C.T., Van Schaick, M.R., Cheema, M.S., Ausió, J., Gowen, B.E., 2019. Spermatogenesis in haploid males of the jewel wasp *Nasonia vitripennis*. Sci Rep 9, 12194. https://doi.org/10.1038/s41598-019-48332-9

Foray, V., Helene, H., Martinez, S., Gibert, P., Desouhant, E., 2013. Occurrence of arrhenotoky and thelytoky in a parasitic wasp Venturia canescens (Hymenoptera: Ichneumonidae): Effect of endosymbionts or existence of two distinct reproductive modes? Eur. J. Entomol. 110, 103–107. https://doi.org/10.14411/eje.2013.014

Giorgini, M., Monti, M.M., Caprio, E., Stouthamer, R., Hunter, M.S., 2009. Feminization and the collapse of haplodiploidy in an asexual parasitoid wasp harboring the bacterial symbiont *Cardinium*. Heredity 102, 365–371. https://doi.org/10.1038/hdy.2008.135

Gloag, R., Remnant, E.J., Oldroyd, B.P., 2019. The frequency of thelytokous parthenogenesis in European-derived *Apis mellifera* virgin queens. Apidologie 50, 295–303. https://doi.org/10.1007/s13592-019-00649-0

Gokhman, V.E., Kuznetsova, V.G., 2018. Parthenogenesis in Hexapoda: holometabolous insects. J Zool Syst Evol Res 56, 23–34. https://doi.org/10.1111/izs.12183

Gu, H., Dorn, S., 2003. Mating system and sex allocation in the gregarious parasitoid *Cotesia glomerata*. Animal Behaviour 66, 259–264. https://doi.org/10.1006/anbe.2003.2185

Hedrick, P.W., Parker, J.D., 1997. Evolutionary Genetics and Genetic Variation of Haplodiploids and X-Linked Genes. Annu. Rev. Ecol. Syst. 28, 55–83. https://doi.org/10.1146/annurev.ecolsys.28.1.55

Heimpel, G.E., de Boer, J.G., 2008. Sex determination in the hymenoptera. Annual review of entomology 53, 209–30. https://doi.org/10.1146/annurev.ento.53.103106.093441

Henter, H.J., 2003. Inbreeding depression and haplodiploidy: experimental measures in a parasitoid and comparisons across diploid and haplodiploid insect taxa. Evolution 57, 1793–1803. https://doi.org/10.1111/i.0014-3820.2003.tb00587.x

Jarosch, A., Stolle, E., Crewe, R.M., Moritz, R.F.A., 2011. Alternative splicing of a single transcription factor drives selfish reproductive behavior in honeybee workers (*Apis mellifera*). Proceedings of the National Academy of Sciences 108, 15282–15287. https://doi.org/10.1073/pnas.1109343108

Kaiser, L., Fernandez-Triana, J., Capdevielle-Dulac, C., Chantre, C., Bodet, M., Kaoula, F., Benoist, R., Calatayud, P.-A., Dupas, S., Herniou, E.A., Jeannette, R., Obonyo, J., Silvain, J.-F., Le Ru, B., 2017. Systematics and biology of *Cotesia typhae sp. n*. (Hymenoptera, Braconidae, Microgastrinae), a potential biological control agent against the noctuid Mediterranean corn borer, Sesamia nonagrioides. ZK 682, 105–136. https://doi.org/10.3897/zookeys.682.13016

Kaiser, L., Le Ru, B.P., Kaoula, F., Paillusson, C., Capdevielle-Dulac, C., Obonyo, J.O., Herniou, E.A., Jancek, S., Branca, A., Calatayud, P., Silvain, J., Dupas, S., 2015. Ongoing ecological speciation in *Cotesia sesamiae*, a biological control agent of cereal stem borers. Evol Appl 8, 807–820. https://doi.org/10.1111/eva.12260

Lattorff, H.M.G., Moritz, R.F.A., Fuchs, S., 2005. A single locus determines thelytokous parthenogenesis of laying honeybee workers (Apis mellifera capensis). Heredity 94, 533–537. https://doi.org/10.1038/sj.hdy.6800654

Leach, I.M., Pannebakker, B.A., Schneider, M.V., Driessen, G., van de Zande, L., Beukeboom, L.W., 2009. Thelytoky in Hymenoptera with Venturia canescens and Leptopilina clavipes as Case Studies, in: Schön, I., Martens, K., Dijk, P. (Eds.), Lost Sex. Springer Netherlands, Dordrecht, pp. 347–375.https://doi.org/10.1007/978-90-481-2770-217

Liegeois, M., Sartori, M., Schwander, T., 2021. Extremely Widespread Parthenogenesis and a Trade-Off Between Alternative Forms of Reproduction in Mayflies (Ephemeroptera). Journal of Heredity 112, 45–57. https://doi.org/10.1093/jhered/esaa027

Little, C.J., Chapuis, M.-P., Blondin, L., Chapuis, E., Jourdan-Pineau, H., 2017. Exploring the relationship between tychoparthenogenesis and inbreeding depression in the Desert Locust, *Schistocerca gregaria*. Ecol Evol 7, 6003–6011. https://doi.org/10.1002/ece3.3103

Liu, Q., Zhou, J., Zhang, C., Ning, S., Duan, L., Dong, H., 2019. Co-occurrence of thelytokous and bisexual Trichogramma dendrolimi Matsumura (Hymenoptera: Trichogrammatidae) in a natural population. Sci Rep 9, 17480. https://doi.org/10.1038/s41598-019-53992-8

Ma, W.-J., Schwander, T., 2017. Patterns and mechanisms in instances of endosymbiont-induced parthenogenesis. J. Evol. Biol. 30, 868–888. https://doi.org/10.1111/jeb.13069

Mackensen, O., 1943. The Occurrence of Parthenogenetic Females in Some Strains of Honeybees. Journal of Economic Entomology 36, 465–467. https://doi.org/10.1093/jee/36.3.465

Meirmans, S., Meirmans, P.G., Kirkendall, L.R., 2012. The Costs Of Sex: Facing Real-world Complexities. The Quarterly Review of Biology 87, 19–40. https://doi.org/10.1086/663945

Mochiah, M.B., Ngi-Song, A.J., Overholt, W.A., Stouthamer, R., 2002. *Wolbachia* infection in *Cotesia sesamiae* (Hymenoptera: Braconidae) causes cytoplasmic incompatibility: implications for biological control. Biological Control 25, 74–80. https://doi.org/10.1016/S1049-9644(02)00045-2

Morgan-Richards, M., Trewick, S.A., 2005. Hybrid origin of a parthenogenetic genus? Molecular Ecology 14, 2133–2142. https://doi.org/10.1111/j.1365-294X.2005.02575.x

Neiman, M., Sharbel, T.F., Schwander, T., 2014. Genetic causes of transitions from sexual reproduction to asexuality in plants and animals. J. Evol. Biol. 27, 1346–1359. https://doi.org/10.1111/jeb.12357

Neumann, P., 2001. Social parasitism by honeybee workers (*Apis mellifera capensis* Escholtz): host finding and resistance of hybrid host colonies. Behavioral Ecology 12, 419–428. https://doi.org/10.1093/beheco/12.4.419

Otto, S.P., 2009. The Evolutionary Enigma of Sex. The American Naturalist 174, S1–S14. https://doi.org/10.1086/599084

Pardo, M.C., López-León, M.D., Cabrero, J., Camacho, J.P.M., 1995. Cytological and developmental analysis of tychoparthenogenesis in Locusta migratoria. Heredity 75, 485–494. https://doi.org/10.1038/hdy.1995.165

Pijls, J.W.A.M., van Steenbergen, H.J., van Alphen, J.J.M., 1996. Asexuality cured: the relations and differences between sexual and asexual *Apoanagyrus diversicornis*. Heredity 76, 506–513. https://doi.org/10.1038/hdy.1996.73

Rabeling, C., Kronauer, D.J.C., 2013. Thelytokous Parthenogenesis in Eusocial Hymenoptera. Annu. Rev. Entomol. 58, 273–292. https://doi.org/10.1146/annurev-ento-120811-153710

Rey, O., Loiseau, A., Facon, B., Foucaud, J., Orivel, J., Cornuet, J.-M., Robert, S., Dobigny, G., Delabie, J.H.C., Mariano, C.D.S.F., Estoup, A., 2011. Meiotic Recombination Dramatically Decreased in Thelytokous Queens of the Little Fire Ant and Their Sexually Produced Workers. Molecular Biology and Evolution 28, 2591–2601. https://doi.org/10.1093/molbev/msr082

Sandrock, C., Vorburger, C., 2011. Single-Locus Recessive Inheritance of Asexual Reproduction in a Parasitoid Wasp. Current Biology 21, 433–437. https://doi.org/10.1016/j.cub.2011.01.070

Schneider, M.V., Beukeboom, L.W., Driessen, G., Lapchin, L., Bernstein, C., Van Alphen, J.J.M., 2002. Geographical distribution and genetic relatedness of sympatrical thelytokous and arrhenotokous populations of the parasitoid Venturia canescens (Hymenoptera): Thelytoky and arrhenotoky in *Venturia canescens*. Journal of Evolutionary Biology 15, 191–200. https://doi.org/10.1046/j.1420-9101.2002.00394.x

Song, Y., Scheu, S., Drossel, B., 2012. The ecological advantage of sexual reproduction in multicellular long-lived organisms: The ecological advantage of sexual reproduction. Journal of Evolutionary Biology 25, 556–565. https://doi.org/10.1111/j.1420-9101.2012.02454.x

Stouthamer, R., Pinto, J.D., Platner, G.R., Luck, R.F., 1990. Taxonomic Status of Thelytokous Forms of *Trichogramma* (Hymenoptera: Trichogrammatidae). Annals of the Entomological Society of America 83, 475–481. https://doi.org/10.1093/aesa/83.3.475

The Tree of Sex Consortium, 2014. Tree of Sex: A database of sexual systems. Sci Data 1, 140015. https://doi.org/10.1038/sdata.2014.15

Tsutsui, Y., Maeto, K., Hamaguchi, K., Isaki, Y., Takami, Y., Naito, T., Miura, K., 2014. Apomictic parthenogenesis in a parasitoid wasp *Meteorus pulchricornis*,uncommon in the haplodiploid order Hymenoptera. Bull. Entomol. Res. 104, 307–313. https://doi.org/10.1017/S0007485314000017

Tucker, K.W., 1958. Automictic parthenogenesis in the Honey bee. Genetics 43, 299–316. https://doi.org/10.1093/genetics/43.3.299

Tvedte, E.S., Logsdon, J.M., Forbes, A.A., 2019. Sex loss in insects: causes of asexuality and consequences for genomes. Current Opinion in Insect Science 31, 77–83. https://doi.org/10.1016/j.cois.2018.11.007

van de Zande, L., Verhulst, E.C., 2014. Genomic Imprinting and Maternal Effect Genes in Haplodiploid Sex Determination. Sex Dev 8, 74–82. https://doi.org/10.1159/000357146

van der Kooi, C.J., Matthey-Doret, C., Schwander, T., 2017. Evolution and comparative ecology of parthenogenesis in haplodiploid arthropods. Evolution Letters 1, 304–316. https://doi.org/10.1002/evl3.30

van Wilgenburg, E., Driessen, G., Beukeboom, L., 2006. Single locus complementary sex determination in Hymenoptera: an “unintelligent” design? Front Zool 3, 1. https://doi.org/10.1186/1742-9994-3-1

van der Kooi, C.J., Schwander, T., 2015. Parthenogenesis: Birth of a New Lineage or Reproductive Accident? Current Biology 25, R659–R661. https://doi.org/10.1016/j.cub.2015.06.055

Verma, S., Ruttner, F., 1983. Cytological analysis of the thelytokous parthenogenesis in the Cape honeybee (*Apis mellifera capensis* Escholtz). Apidologie 14, 41–57. https://doi.org/10.1051/apido:19830104

Vershinina, A.O., Kuznetsova, V.G., 2016. Parthenogenesis in Hexapoda: Entognatha and non-holometabolous insects. J Zoolog Syst Evol Res 54, 257–268. https://doi.org/10.1111/jzs.12141

Vorburger, C., 2014. Thelytoky and Sex Determination in the Hymenoptera: Mutual Constraints. Sex Dev 8, 50–58. https://doi.org/10.1159/000356508

Weeks, E.N., Birkett, M.A., Cameron, M.M., Pickett, J.A., Logan, J.G., 2011. Semiochemicals of the common bed bug, *Cimex lectularius L*. (Hemiptera: Cimicidae), and their potential for use in monitoring and control. Pest. Manag. Sci. 67, 10–20. https://doi.org/10.1002/ps.2024

Whiting, P.W., 1945. The Evolution of Male Haploidy. The Quarterly Review of Biology 20, 231–260. https://doi.org/10.1086/394884

Zchori-Fein, E., Perlman, S.J., 2004. Distribution of the bacterial symbiont Cardinium in arthropods: Molecular Ecology 13, 2009–2016. https://doi.org/10.1111/j.1365-294X.2004.02203.x

Zhou, Y., Gu, H., Dorn, S., 2007. Effects of inbreeding on fitness components of *Cotesia glomerata*, a parasitoid wasp with single-locus complementary sex determination (sl-CSD). Biological Control 40, 273–279. https://doi.org/10.1016/j.biocontrol.2006.11.002

