## supplementary_data for "Spontaneous parthenogenesis in the parasitoid wasp *Cotesia typhae*: low frequency anomaly or evolving process?"

| SNP marker | position (cM) | chromosome | K4M1-F2L-1 | K4M1-F2O-1 | K4M1-F2O-2 | K4M1-F2O-3 | K4M1-F2O-4 | K5M1-F2C-1 | K5M1-F2G-1 | M5K2-F2C-1 | M5K5-F2M-1 |
|---|---|---|---|---|---|---|---|---|---|---|---|
| 1471_nov | 0 | 1 | MM | KM | KK | KK | KM | KM | KM | KK | KM |
| 5590_nov | 37.5 | 1 | KM | KM | KM | KM | KM | KM | KM | KM | KM |
| 362_nov | 88.9 | 1 | KM | KM | KK | KM | KM | KM | KM | KM | KM? |
| **chromosome length** | 97.8 | 1 | | | | | | | | | |
| 18875_nov | 3.3 | 2 | MM | KM | KK | KK | KM | KK | KM | MM | KM |
| 8225_nov | 17.6 | 2 | ? | KM | KK | KK | KM | KK | KM | MM | KM |
| 10640_nov | 28.6 | 2 | KM? | KM | KK | KM | KM | KK | KM | MM | KM |
| 32759_nov | 36.3 | 2 | KM | KM | KM | KM | KM | KM | KM | KM | KM |
| 1975_nov | 50.3 | 2 | KM | KM | KM | KM | KM | KM | KM | KM | KM |
| 13818_nov | 60.4 | 2 | KM | KM | KM | KM | KM | KM | KM | KM | KM |
| 11549_nov | 69.2 | 2 | KM | KM | KM | KM | KM | KM | KM | KM | KM |
| 30693_nov | 84.6 | 2 | KM | KM | KK | KM | KM | KK | KM | KM | KM |
| 16491_nov | 97.9 | 2 | KM | KM | KK | KM | KM | KK | KM | KM | KM |
| 8884_nov | 106.7 | 2 | KM | KM | KM | KM | KK | KK | KM | KM | KM |
| 2333_nov | 119.9 | 2 | KM | KM | KM | MM | KK | KK | KM | KK | KM |
| 186_nov | 136.4 | 2 | KM | KM | KM | MM | KK | KK | KM | KK | KM |
| **chromosome length** | 136.4 | 2 | | | | | | | | | |
| 14555_nov | 1.1 | 3 | KM | KM | KM | KM | MM | KM | KM | MM | KM |
| 27134_nov | 38.6 | 3 | KM | KM | KK | KM | KM | KM | KM | KM | KM |
| 21770_nov | 86.1 | 3 | ? | KM | KK | KM | KM | KM | KM | KM | KM |
| **chromosome length** | 89.4 | 3 | | | | | | | | | |
| 19620_nov | 19.8 | 4 | KM | KM | MM | KM | KM | MM | KM | KM | KM |
| 20912_ref | 111.4 | 4 | ? | KM | KM | KM | MM | KM | KM | MM | KM |
| **chromosome length** | 126.8 | 4 | | | | | | | | | |
| 8760_nov | 2.2 | 5 | KM | KM | KK | KM | KM | KK | KM | KK | KM |
| 24361_nov | 112.6 | 5 | KM | KM | KM | KM | KM | KM | KM | MM | KM |
| **chromosome length** | 117 | 5 | | | | | | | | | |
| 9613_nov | 2.2 | 6 | KK | KM | KM | KM | KM | KM | KM | KM | KM |
| 10874_nov | 9.9 | 6 | KM | KM | KM | KM | KM | KM | KM | KM | KM |
| 24116_nov | 23.1 | 6 | KK | KM | KK | KM | KM | KM | KM | KM | KM |
| 26551_nov | 30.8 | 6 | KM | KM | KK | KM | KM | KM | KM | KM | KM |
| 20921_ref | 65.9 | 6 | KM | KM | KK | KM | KM | KM | KM | KM | KM |
| 252_nov | 76.9 | 6 | KM | KM | KK | KM | KM | KM | KM | KM | KM |
| 1191_nov | 85.7 | 6 | KM | KM | KM | KM | KM | KM | KM | KM | KM |
| 21110_nov | 94.5 | 6 | KM | KM | KM | KM | KM | KM | KM | KM | KM |
| 32954_nov | 120.9 | 6 | KM | KM | KM | KM | KM | KM | KM | KM | KM |
| 9911_nov | 129.7 | 6 | KM | KM | KM | KM | KM | KM | KM | KM | KM |
| 25277_nov | 157.5 | 6 | KM | KM | KK | KM | MM | KK | KM | KM | KM |
| 13687_nov | 166.3 | 6 | KM | KM | KK | KM | MM | KK | KM | KM | KM? |
| **chromosome length** | 166.3 | 6 | | | | | | | | | |
| 26354_nov | 3.3 | 7 | KM | KM | MM | KK | KM | KM | KM | KM | KM |
| 28179_nov | 16.5 | 7 | KM | KM | KM | KK | KM | KM | KM | KM | KM |
| 29710_nov | 39.6 | 7 | KM | KM | KM | KK | KM | KM | KM | KM | KM |
| 17508_nov | 59.4 | 7 | KM | KM | KM | KK | KM | KM | KM | KM | KM |
| 10925_nov | 86.9 | 7 | KM | KM | KM | KM | KM | KM | KM | KM | KM |
| 23654_ref | 95.7 | 7 | KM | KM | KM | KM | KM | KM | KM | KM | KM |
| 2013_nov | 112.2 | 7 | KM | KM | KM | KM | KM | KM | KM | KM | KM |
| 13126_nov | 123.2 | 7 | KM | KM | KK | KM | KM | KM | KM | KM | KM |
| 25998_nov | 133.1 | 7 | KM | KM | KK | KM | KM | KM | KM | KM | KM |
| 27068_nov | 144.1 | 7 | KM | KM | KK | KM | KM | KM | KM | KM | KM |
| 10217_nov | 159.5 | 7 | KM | KM | KK | KM | MM | KM | KM | MM | KM |
| 6713_nov | 167.2 | 7 | MM? | KM | KK | KM | MM | KM | KM | MM | KM |
| **chromosome length** | 170.5 | 7 | | | | | | | | | |
| 17926_nov | 0 | 8 | KM | KM | KK | KM | MM | KM | KM | KK | KM |
| 23207_nov | 13.2 | 8 | KM | KM | KK | KM | MM | KM | KM | KK | KM |
| 8454_nov | 24.2 | 8 | KM | KM | KK | KM | KM | KM | KM | KK | KM |
| 32305_nov | 37.4 | 8 | KM | KM | KM | KM | KM | KM | KM | KM | KM |
| 14447_nov | 46.2 | 8 | KM | KM | KM | KM | KM | KM | KM | KM | KM |
| 20609_nov | 57.2 | 8 | KM | KM | KM | KM | KM | KM | KM | KM | KM |
| 32961_nov | 84.7 | 8 | KM | KM | KM | KM | KM | KK | KM | KM | KM |
| 9051_nov | 94.6 | 8 | MM | KM | KM | KM | KM | KK | KM | KM | KM |
| 9503_nov | 116.6 | 8 | MM | KM | KM | KM | KM | KK | KM | KM | KM |
| 11486_nov | 136.4 | 8 | KM | KM | KM | KM | KK | KK | KM | KM | KM |
| **chromosome length** | 145.2 | 8 | | | | | | | | | |
| 20661_nov | 4.5 | 9 | ? | KM | KM | KK | MM | KM | KM | KM | KM |
| 18209_nov | 41.9 | 9 | KM | KM | KM | KK | KM | KM | KM | KM | KM |
| 5560_nov | 69.4 | 9 | KM | KM | KM | KM | KM | KM | KM | KM | KM |
| 19000_nov | 107.1 | 9 | KM | KM | KM | KM | KM | KM | KM | KM | KM |
| **chromosome length** | 107.1 | 9 | | | | | | | | | |
| 10511_nov | 1.1 | 10 | KM | KM | MM | KM | MM | KK | KM | MM | KM |
| 10299_nov | 47.6 | 10 | KM | KM | KM | KM | MM | KM | KM | KM | KM |
| 3191_nov | 109.6 | 10 | KM | KM | MM | KM | KM | KM | KM | KM | KM |
| **chromosome length** | 144.2 | 10 | | | | | | | | | |

Supplementary Table 1: Genotypes of the 9 parthenogenetic daughters of virgin heterozygous females at all the 63 SNP markers used. The markers are ordered according to their position on the chromosomes and the colours correspond to the 3 different possible genotypes: yellow for Makindu homozygote, green for heterozygote, and blue for Kobodo homozygote. A change of colour in a parthenogenetic daughter's column indicates a recombination event between the two successive markers. Grey cells correspond to uncertain genotypes.

| SNP marker | forward primer (5'-3') | reverse primer (5'-3') | Kobodo allele | Makindu allele | genotyping method |
|---|---|---|---|---|---|
| 1471_nov | TCCGTATAAGGTAAAAGTCCCCA | TTGCCGAATCCTTCTGTGGT | A | G | HRM |
| 5590_nov | AACCAGTTGCTGCACGAC | AGTGCTAGCTCTTTCGGCA | T | C | HRM |
| 362_nov | CTACGCCCAAGTGCATCAAC | AAAGCGCATGATGTCCCTCT | T | C | HRM |
| 18875_nov | TTGACGAAACCTTTAGACGGG | GCAGTGCAGACAATATTTACCGG | C | T | HRM |
| 8225_nov | Makindu specific : CTGGTTTTTCTGGTATTTCACCTT<br>Kobodo specific : CTGGTTTTTCTGGTATTTCACCTC | CGTTTAACAGCAAGTGGCTTA | C | T | allele specific PCR |
| 10640_nov | CGGATGCTGCAGAGGTTG | CAAGTTTCCCAGTGCCATCG | T | C | HRM |
| 32759_nov | ATACATCAATGGCGCATGGG | TCGAGTACTTCTTGTCACAACA | T | C | HRM |
| 1975_nov | ACACTGCGCTCCTAAGAACA | GACTCGGTCTCACTGCAGT | G | C | HRM |
| 13818_nov | CAGCGTTTGTCTACTGCAGA | TGCATCTATATCACGCTGTTCA | C | T | HRM |
| 11549_nov | AGACTCAAATGACAGCTGCAGA | CTCCTCCGCGGTGATAACTG | T | C | HRM |
| 30693_nov | TGCCTTTGTCTTTGTCTTTGTCT | CGTTCTGCAGTTCCAACACC | C | T | HRM |
| 16491_nov | GTTCGGTCACAGTGATGCCT | TGTAGCAACAGCCTCACCT | T | C | HRM |
| 8884_nov | TGATGTCTTCGAGCTGGTCT | AGTGCACTTCTACTCCACGC | A | G | HRM |
| 2333_nov | AGTTTCATTTATCGGCTGCAGG | CGCTGTGTAGTTTGTGCTTCA | A | G | HRM |
| 186_nov | CGTGCCATCTGTACTTCTCCA | TGGAAAGAAATTAGATCGGCGGA | A | G | HRM |
| 14555_nov | CCCTCTCGCTGCTGAATAGG | GGACCACCTTTACAGCTGCA | G | T | HRM |
| 27134_nov | AGCAGACTGAGGAATTGGATGG | GCAACGTCTCCACTCTCACA | G | A | HRM |
| 21770_nov | TCTGGGCTATAGGAATTCAA | Makindu specific : CAGTATTGTCAGCTGAGGA<br>Kobodo specific : CAGTATTGTCAGCTGAGGG | C | T | allele specific PCR |
| 19620_nov | AAACTCACGGTGCTGCAGTA | ACGCTGGCATTGGAGGAATT | G | A | HRM |
| 20912_ref | GCCATAGCGAGGTTAACGG | ATGCAGTACCTTACGCCGAG | A | G | HRM |
| 8760_nov | GACCACCGACAATCCCATCA | AAGGGCATCTGAAGGTGGTC | T | C | HRM |
| 24361_nov | AGTCAGCCTGCAGTTTCCAA | CGTACAGCATCACACGGAGG | T | C | HRM |
| 9613_nov | TCACATCCTTTCTGACGACGT | CAGCGAGAAGGACTGGTGC | A | G | HRM |
| 10874_nov | CACAAAGGACCCGCATTTCA | TCCGGCTTATTCTCCCTGTT | G | A | HRM |
| 24116_nov | GTGCTTCGGTGCCCTTAAG | ACTGCAGTATAGGTGAGTTTGGT | G | A | HRM |
| 26551_nov | GGGGAGGGGCGGAATTACTA | TCACACACAATGACGGTACA | C | T | HRM |
| 20921_ref | GGACTCGGTTGTGTGAACTCT | TGAGACTCACCGTTACCGTC | T | G | HRM |
| 252_nov | GAGTGGCGCTAGTGAAGATCT | AGGTTTTATGGAATTGCTGCAGT | G | A | HRM |
| 1191_nov | ACTGTTTCTCCTGTTCTTTCCT | ACCCTTCATTTCCAGAGCCG | A | G | HRM |
| 21110_nov | GGAACTTCCCGCAATATACGA | AGTGGTGTTTGTGGGTTTGA | G | T | HRM |
| 32954_nov | AACAATTTGGCACGTCAGGT | TGGACTTTACTCTCGTTTGCTCA | C | T | HRM |
| 9911_nov | ACAAAACACAAACTGCAGGACT | ATGCACGCGGAACTTGTCTA | G | A | HRM |
| 25277_nov | ACACCGTAGATCTCAGATCGT | GCAAAGATCGTGAAAGTGGCA | T | C | HRM |
| 13687_nov | GAAGCAGAGTTCCGACGAGT | AAGGCGAACTGGAACATTTG | T | C | HRM |
| 26354_nov | TCGCGTGTTTATCTGTCGGA | TGCCTCTTTTACTTCCTGCAGA | A | G | HRM |
| 28179_nov | TGGCTCTTAATGTTTGCGACG | ACTTAATACACGGCATCTGGAA | A | G | HRM |
| 29710_nov | AGCATTACTACTGGGACCGG | AATTGTGATGCTGGCTGGGA | A | G | HRM |
| 17508_nov | GTCTCCACTTTGCATGCATG | GGATGACTAATGATCTCTGCAGG | G | A | HRM |
| 10925_nov | AATTAAACCTCCACATCTGCGT | CGCAAACAGGACCATTAAACGT | C | T | HRM |

| | | | | | |
|---|---|---|---|---|---|
| 23654_ref | CCCCTGACGGCAACATACAT | GCGACGGTTACTGCTCTGAT | G | A | HRM |
| 2013_nov | CGAACGTTTTGAGCGAGCAT | TGCTTCCAAGTACATCGGAGG | G | A | HRM |
| 13126_nov | CTGCAGGCAAGAACATTGTT | GCAAGCATGGTTATTGAGGACA | C | T | HRM |
| 25998_nov | AACACTGTCCAAACTTCCTGT | TGGGTACTCAAAACTGCAGCA | G | T | HRM |
| 27068_nov | CCTCCTGCAGCTCTTCTTCC | CCGTTGTTAGTTGTTGCTTTGG | A | G | HRM |
| 10217_nov | CGAGCTTTCATTGTCTACTGGC | CGTGTGAACAGAACAGTATCGG | G | A | HRM |
| 6713_nov | TCCGTGTGAGTGCTTTCGTT | TGAGTCTGCAGTATCTAACGACA | C | T | HRM |
| 17926_nov | GCCGGGCAATAATCCAGAAG | AGAGCTTGGCGGTGATGG | G | A | HRM |
| 23207_nov | CTTGATCAAGCAGGCGTTG | GGCACTGCAGGCTATGTTAA | G | A | HRM |
| 8454_nov | GAAGACCAGCGCCGACTTTA | GCGATGAAACTTCTAGACCTGC | C | T | HRM |
| 32305_nov | TCAACGACAACTGCAGGTACA | GACGTCGTACCAGATGCCG | G | T | HRM |
| 14447_nov | TGGGATGATCTGGGTCTGTG | CGCTGCAGAAGATCGTCGAT | C | T | HRM |
| 20609_nov | ACAGGTAGGCGCGATATTCC | CCGCGTCTACTCAGTACCAC | C | T | HRM |
| 32961_nov | ATTTTCGACTGCTGGGGAGG | CAATGTATGCTGCAGAGTCACA | T | C | HRM |
| 9051_nov | AACAGAACAGGAATAGTAGCAGT | CGGTAATATTGATCGCATCTCCG | CA | TT | HRM |
| 9503_nov | ATTCCAGTGCTGCAGTCCAA | ATTTGATGCCGCGAGACTCA | C | T | HRM |
| 11486_nov | TCCCCAGGCTCTGCAGTC | ACAACTGCATACAAGAGTGATGG | C | T | HRM |
| 20661_nov | CGCTATCGGCTTATTCTCCA | AAGCACCCAAATTGTCCTTG | C | T | HRM |
| 18209_nov | CAGCAGTATCCGCCCAGAAT | ATTGACACCTGGGACTGCAG | T | C | HRM |
| 5560_nov | TCTTGGAACTGACGCTGACC | TCGCTGGAACGTCAAGTCTC | A | G | HRM |
| 19000_nov | CGTTCAACTAGGCGAGCAGA | CTGCGGGATTCATGGTGATAA | A | G | HRM |
| 10511_nov | TGTGACCTCTGCAGACCATG | ACGACGAGTCAAAGCAAAGTG | G | A | HRM |
| 10299_nov | TCACCGTTAATTCGTCCCTCA | GCTTGCTTGGTTCCTCTCGA | C | T | HRM |
| 3191_nov | AACCGCTTTTACCCCGAGAG | CGACGGAATTGGCGACCT | C | T | HRM |

Supplementary Table 2: Primers’ sequences for the 63 SNP markers used for genotyping. The Kobodo and Makindu reference alleles are also indicated for each SNP.

| Micro-organism (gene) | Primers | Sequence (5'-3') | Tm | Size (bp) | Reference |
|---|---|---|---|---|---|
| Wolbachia (gltA) | WgltAF1 | TACGATCCAGGGTTTGTTTCTAC | 55°C | 750 | Casiraghi et al., 2005 |
| | WgltARev1 | CTCATTAGCTCCACCGTGTG | | | |
| Bacteroidetes (16S) | Ch-F | TACTGTAAGAATAAGCACGGC | 57°C | 900 | Zchori-Fein and Perlman, 2004 |
| | Ch-R | GTGGATCACTTAACGCTTTCG | | | |
| Cardinium (16S) | CLO-F | GGAACCTTACCTGGGCTAGAATGTATT | 56°C | 500 | Gotoh et al., 2007 |
| | CLO-R | GCCACTGTCTTCAAGCTCTACCAAC | | | |
| Rickettsia (16S) | Rb-F | GCTCAGAACGAACGCTATC | 56°C | 900 | Gottlieb et al., 2006 |
| | Rb-R | GAAGGAAAGCATCTCTGC | | | |
| Spiroplasma (16S) | Spixo-F | TTAGGGGCTCAACCCCTAACC | 52°C | 810 | Duron et al., 2008 |
| | Spixo-R | TCTGGCATTGCCAACTCTC | | | |
| Arsenophonus (23S) | Ars23s-F | CGTTTGATGAATTCATAGTCAAA | 52°C | 650 | Thao and Baumann, 2004 |
| | Ars23s-R | GGTCCTCCAGTTAGTGTTACCCAAC | | | |
| Microsporidia (16S) | V1-F | CACCAGGTTGATTCT | 57°C | 1300 | Weiss et al., 1994 |
| | 1492-R | GGTTACCTTGTTACGACTT | | | Weiss and Vossbrinck, |
| | V1-F | CACCAGGTTGATTCT | 63°C | 450 | 1998 |
| | 530-R | CCGCGGCTGCTGGCAC | | | Baker et al., 1995 |

Supplementary Table 3: List of the micro-organisms whose presence was tested in *C. typhae* Makindu females, by PCR with specific primers.
